# Dopamine Synthesis Capacity and Tau Pathology Differentially Modulate Rest-to-Task Functional Network Dynamics

**DOI:** 10.64898/2026.11.04.764247

**Authors:** Thomas M. Morin, Hsiang-Yu Chen, Shaian Aghasoltan, Ming Hsu, William J. Jagust, Anne S. Berry

## Abstract

Optimal cognition relies on large-scale functional brain networks which balance segregation at rest with selective integration during task performance. In aging, these network dynamics are altered. We examined two prominent features of brain aging: dysregulated dopamine function and Alzheimer’s disease (AD)-related pathology to examine impacts on brain network dynamics. Young adults and cognitively unimpaired older adults completed resting-state and task-based fMRI and PET imaging of dopamine synthesis capacity ([^18^F]fluoro-m-L-tyrosine). Older adults were classified as β-amyloid negative/positive using [^11^C]Pittsburgh Compound-B PET (PiB-or PiB+) and underwent tau-PET scanning ([^18^F]flortaucipir). Greater rest-to-task network stability and task-related network segregation were associated with better subsequent memory. Compared to young adults, PiB+ older adults showed reduced network stability and both PiB+ and PiB-older adults showed reduced segregation during rest. Higher ventral striatal dopamine synthesis capacity was associated with greater stability within associative networks in young adults, but this relationship was attenuated in older adults and absent in PiB+ participants. In PiB+ older adults, higher tau burden was associated with a task-induced collapse of network segregation across large-scale networks. These findings reveal dissociable influences of dopamine and tau on rest-to-task network organization, highlighting the value of integrating neuromodulatory and pathological markers in aging research.

**Author Summary:** As we age, our brains become less efficient at balancing network segregation at rest with selective integration during task states. In this study, we combine brain imaging methods to examine how dopamine, a key neuromodulator, and tau pathology, a hallmark of Alzheimer’s disease, influence large-scale brain network dynamics. We show that dopamine supports stable and well-organized brain networks while tau pathology disrupts this organization during a cognitive task. These opposing influences help explain why some individuals maintain cognitive resilience while others show early network dysfunction, providing new insight into mechanisms of cognitive aging and age-related pathology.

## Introduction

The human brain responds to varying cognitive demands by balancing functional segregation with selective integration of large-scale networks. During rest, large-scale brain networks tend to exhibit stable, highly segregated connectivity patterns, reflecting specialized processing within functional systems. Task engagement, in contrast, induces selective integration between networks to support cognition, while preserving a stable core network architecture (Gratton et al., 2016, 2018; D. H. Schultz & Cole, 2016; Shine et al., 2016). These dynamics can be quantified by measuring rest-to-task network stability (the similarity of connectivity patterns across states), and network segregation (the relative balance of within-and between-network connectivity). In young adults, greater rest-to-task network stability and segregation predict better cognition (including memory, reasoning, and executive functioning), whereas larger rest-to-task shifts generally predict worse performance (Ferguson et al., 2017; Hilger et al., 2020; D. H. Schultz & Cole, 2016; Thiele et al., 2022). Put simply, the maintenance of a stable, segregated network across resting and task states is increasingly recognized as a hallmark of healthy cognition across a range of cognitive domains (Chan et al., 2014; Krienen et al., 2014; D. H. Schultz & Cole, 2016; Wang et al., 2021).

The maintenance of stable, segregated networks across rest and task states appears to be reduced in older age. Analyses of large, multi-center fMRI studies have demonstrated that normal aging is characterized by reduced network segregation at rest (Betzel et al., 2014; Chan et al., 2014). Longitudinal evidence suggests that age-related reductions in resting-state network segregation are associated with poorer cognitive outcomes (Chan et al., 2021), and reduced gains following cognitive training (Gallen et al., 2016; Gallen & D’Esposito, 2019). Prior work has also proposed that age-related reductions in resting-state segregation may incite unstable network dynamics during rest-to-task state transitions (Chan et al., 2017; Cole et al., 2016). Emerging research focused on the dynamics of large networks finds lower rest-to-task network stability in older age, which has been associated with increased errors on motor (Monteiro et al., 2019), episodic memory (Cassady et al., 2023), and working memory tasks (Yue et al., 2024). These age-related alterations in network dynamics are posited to contribute to cognitive decline, yet the biological mechanisms underlying them are not completely understood.

In young adults, positron emission tomography (PET) has linked endogenous dopamine measures with individual differences in network coupling among default mode, dorsal attention and frontoparietal control functional networks during rest (Dang et al., 2012), and stronger within-network connectivity during rest and task (Roffman et al., 2017). Beyond functional connectivity strength, however, no study has directly tested whether dopamine PET measures relate to network segregation or rest-to-task stability. Converging work using pharmacology, pupillometry-fMRI, genetic variation, and computational modeling links dopaminergic signaling to large-scale segregation, flexibility, and stability (Braun et al., 2021; Garrett et al., 2015; Hansen et al., 2022; Shine, 2023; Shine et al., 2021). In particular, computational models suggest that neuromodulators influence global network organization via diffuse projections throughout cortex that modulate neural gain (defined as the signal-to-noise ratio of afferent inputs projecting from subcortical neuromodulatory nuclei) (Shine, 2023; Shine et al., 2018). Though the dopamine system substantially changes during aging (Karalija et al., 2022; Karrer et al., 2017; Lundgren et al., 2025; Nevalainen et al., 2015), there is relatively little research investigating the nature of dopaminergic modulation of large-scale functional network dynamics across the lifespan.

Aging is accompanied by the accumulation of neuropathology including amyloid-β, and tau pathology. The development and spread of tau pathology appears to follow patterns of functional connectivity, suggesting network architecture may shape how pathology spreads (Adams et al., 2019; Franzmeier et al., 2020; Vogel et al., 2020). Consistent with this view, recent studies report that greater tau burden is associated with altered task-related network reconfiguration and reduced network segregation in cognitively unimpaired older adults with elevated Aβ (PiB+) (Adams et al., 2023; Cassady et al., 2023). Together, these findings indicate that tau pathology may contribute to network instability, manifesting as excessive network integration and less efficient transitions between rest and task states, particularly in the presence of Aβ.

We hypothesized that dopaminergic modulation and tau pathology may exert opposing influences on large-scale network dynamics. Whereas dopamine is thought to stabilize network configuration and preserve functional segregation, tau pathology may promote aberrant integration and task-induced collapse of network boundaries. Understanding how these competing forces interact is critical for explaining heterogeneity in cognitive aging trajectories. In the present study, we integrate multimodal neuroimaging to examine how dopamine synthesis capacity (measured with [^18^F]fluoro-m-L-tyrosine (FMT) PET) and tau pathology (measured with [^18^F]flortaucipir (FTP) PET) jointly shape rest-to-task network stability (measured with fMRI) in both young and older adults. To quantify network stability, we compared network organization during the memory-encoding portion of a task with resting-state organization measured both before and after the task. Because network stability may depend on task demands, we used a subsequent memory paradigm that involved incidental encoding during a reward learning task (Morin et al., 2026), to test whether individual differences in dopamine synthesis capacity and tau pathology are associated with network stability in a dopamine-relevant context.

By directly linking PET-derived measures of dopamine synthesis and tau pathology with fMRI-based network dynamics, this work provides a unique opportunity to dissociate neuromodulatory and pathological contributions to large-scale brain organization. We test the hypothesis that dopaminergic function supports stable and segregated network architecture, particularly within associative networks, whereas tau pathology undermines segregation during task engagement in the context of high cerebral amyloid-β load. Together, these analyses aim to clarify how molecular-level processes shape network-level dynamics, with implications for understanding cognitive aging and the earliest stages of AD.

## Results

### Experimental Overview

Participants were young adults and cognitively unimpaired older adults from the Berkeley Aging Cohort Study (demographics in **Table 1**). During a single MRI session, participants completed a pre-task resting-state scan, four runs of a reward-learning task scan, and a post-task resting-state scan. Recognition memory for task stimuli was assessed ∼30 minutes later outside of the scanner. In the task, participants learned, via trial-by-trial monetary feedback, to predict whether houses shown with different realtors would be sold for a profit, break even, or sell at a loss. Specific task details and univariate results have been previously described (Morin et al., 2026).The subsequent memory test probed recognition for the house images encountered during the task.

**Table 1.**
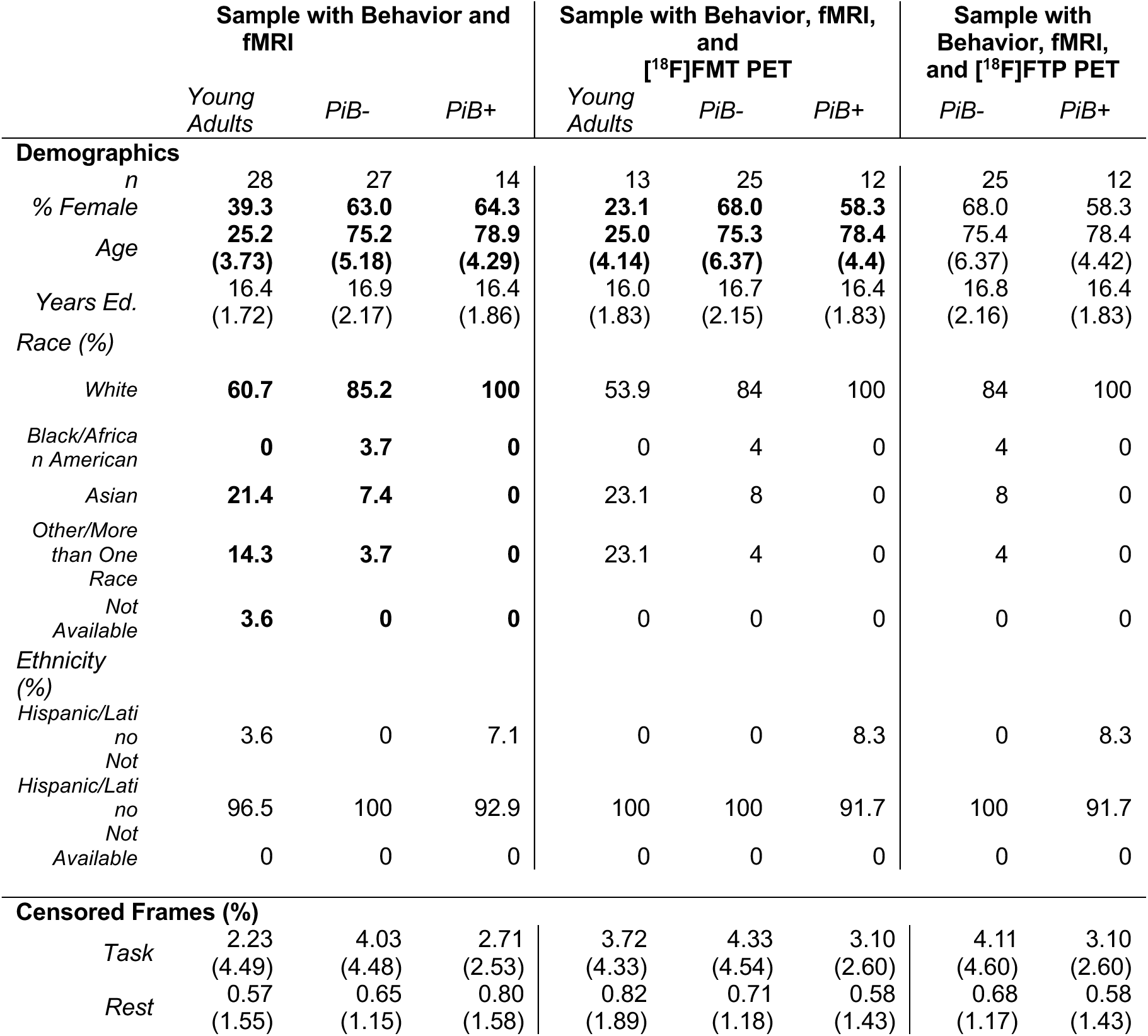
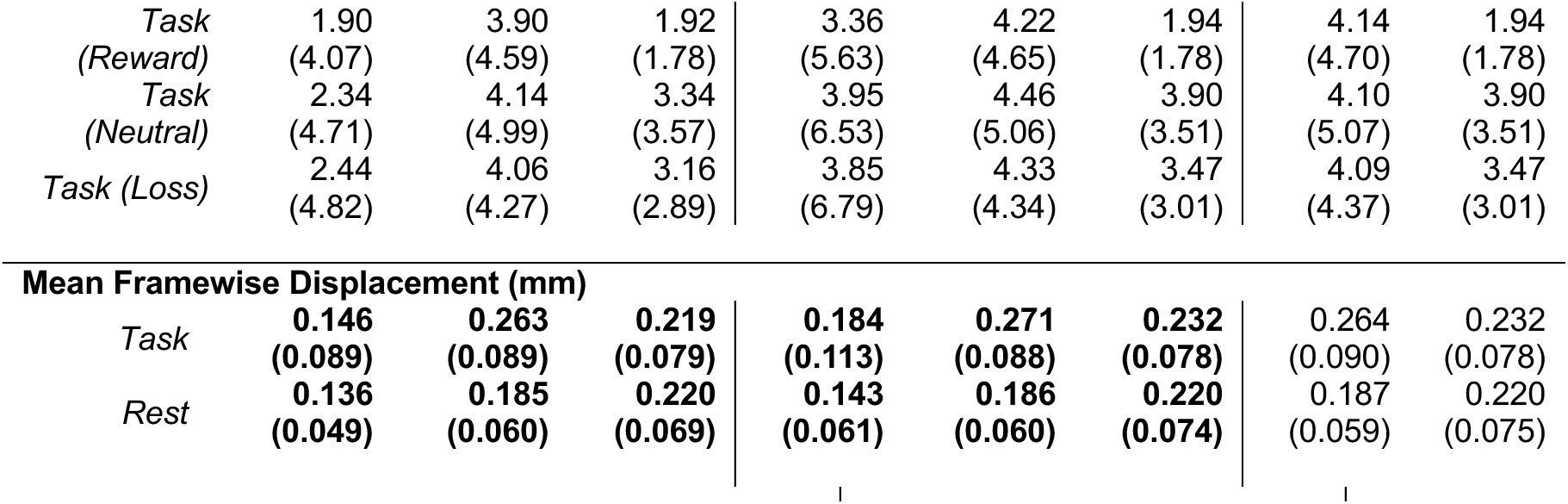
Sample Demographics. Participant demographics are displayed for participants who passed data quality control and were included in the analyses. Mean and (Standard Deviation) are presented. % Censored Frames represents the percentage of frames that were censored from the fMRI data due head motion (framewise displacement > 0.5mm and/or DVARS > 5mm). Mean framewise displacement (mm) reflects the mean frame-to-frame head motion in the final fMRI data after censoring for high motion and ensuring equal quantities of data across conditions. Bold rows indicate a significant group difference in a metric using ANOVA (for three groups) or a T-test (for two groups).

|  | **Sample with Behavior and fMRI** |  |  | **Sample with Behavior, fMRI, and [<sup>18</sup>F]FMT PET** |  |  | **Sample with Behavior, fMRI, and [<sup>18</sup>F]FTP PET** |  |
|---|---|---|---|---|---|---|---|---|
|  | *Young Adults* | *PiB-* | *PiB+* | *Young Adults* | *PiB-* | *PiB+* | *PiB-* | *PiB+* |
| **Demographics** |  |  |  |  |  |  |  |  |
| *n* | 28 | 27 | 14 | 13 | 25 | 12 | 25 | 12 |
| *% Female* | **39.3** | **63.0** | **64.3** | **23.1** | **68.0** | **58.3** | 68.0 | 58.3 |
| *Age* | **25.2<br>(3.73)** | **75.2<br>(5.18)** | **78.9<br>(4.29)** | **25.0<br>(4.14)** | **75.3<br>(6.37)** | **78.4<br>(4.4)** | 75.4<br>(6.37) | 78.4<br>(4.42) |
| *Years Ed.* | 16.4<br>(1.72) | 16.9<br>(2.17) | 16.4<br>(1.86) | 16.0<br>(1.83) | 16.7<br>(2.15) | 16.4<br>(1.83) | 16.8<br>(2.16) | 16.4<br>(1.83) |
| *Race (%)* |  |  |  |  |  |  |  |  |
| *White* | **60.7** | **85.2** | **100** | 53.9 | 84 | 100 | 84 | 100 |
| *Black/African American* | **0** | **3.7** | **0** | 0 | 4 | 0 | 4 | 0 |
| *Asian* | **21.4** | **7.4** | **0** | 23.1 | 8 | 0 | 8 | 0 |
| *Other/More than One Race* | **14.3** | **3.7** | **0** | 23.1 | 4 | 0 | 4 | 0 |
| *Not Available* | **3.6** | **0** | **0** | 0 | 0 | 0 | 0 | 0 |
| *Ethnicity (%)* |  |  |  |  |  |  |  |  |
| *Hispanic/Latino* | 3.6 | 0 | 7.1 | 0 | 0 | 8.3 | 0 | 8.3 |
| *Not Hispanic/Latino* | 96.5 | 100 | 92.9 | 100 | 100 | 91.7 | 100 | 91.7 |
| *Not Available* | 0 | 0 | 0 | 0 | 0 | 0 | 0 | 0 |
| **Censored Frames (%)** |  |  |  |  |  |  |  |  |
| *Task* | 2.23<br>(4.49) | 4.03<br>(4.48) | 2.71<br>(2.53) | 3.72<br>(4.33) | 4.33<br>(4.54) | 3.10<br>(2.60) | 4.11<br>(4.60) | 3.10<br>(2.60) |
| *Rest* | 0.57<br>(1.55) | 0.65<br>(1.15) | 0.80<br>(1.58) | 0.82<br>(1.89) | 0.71<br>(1.18) | 0.58<br>(1.43) | 0.68<br>(1.17) | 0.58<br>(1.43) |
| *Task (Reward)* | 1.90<br>(4.07) | 3.90<br>(4.59) | 1.92<br>(1.78) | 3.36<br>(5.63) | 4.22<br>(4.65) | 1.94<br>(1.78) | 4.14<br>(4.70) | 1.94<br>(1.78) |
| *Task (Neutral)* | 2.34<br>(4.71) | 4.14<br>(4.99) | 3.34<br>(3.57) | 3.95<br>(6.53) | 4.46<br>(5.06) | 3.90<br>(3.51) | 4.10<br>(5.07) | 3.90<br>(3.51) |
| *Task (Loss)* | 2.44<br>(4.82) | 4.06<br>(4.27) | 3.16<br>(2.89) | 3.85<br>(6.79) | 4.33<br>(4.34) | 3.47<br>(3.01) | 4.09<br>(4.37) | 3.47<br>(3.01) |
| **Mean Framewise Displacement (mm)** |  |  |  |  |  |  |  |  |
| *Task* | **0.146<br>(0.089)** | **0.263<br>(0.089)** | **0.219<br>(0.079)** | **0.184<br>(0.113)** | **0.271<br>(0.088)** | **0.232<br>(0.078)** | 0.264<br>(0.090) | 0.232<br>(0.078) |
| *Rest* | **0.136<br>(0.049)** | **0.185<br>(0.060)** | **0.220<br>(0.069)** | **0.143<br>(0.061)** | **0.186<br>(0.060)** | **0.220<br>(0.074)** | 0.187<br>(0.059) | 0.220<br>(0.075) |

A subset of participants completed PET imaging of dopamine synthesis capacity using [^18^F]fluoro-m-L-tyrosine (FMT). In a subset of older adults, tau burden was quantified with [^18^F]flortaucipir (FTP), and amyloid-β burden was assessed with [^11^C]Pittsburgh Compound B (PiB). Older adults were classified as amyloid-negative (PiB−) or amyloid-positive (PiB+) using a global DVR cutoff (gray matter/cerebellar gray = 1.065; Villeneuve et al., 2015). One older adult participant without PiB data was assumed PiB-. Functional connectivity networks were estimated for each rest run and for each task condition (see **Figure 1** and **Methods** for full details). Analyses tested whether individual differences in dopamine synthesis capacity and tau pathology relate to large-scale network dynamics (stability and segregation), and whether these network dynamics related to subsequent recognition memory.

**Figure 1.**
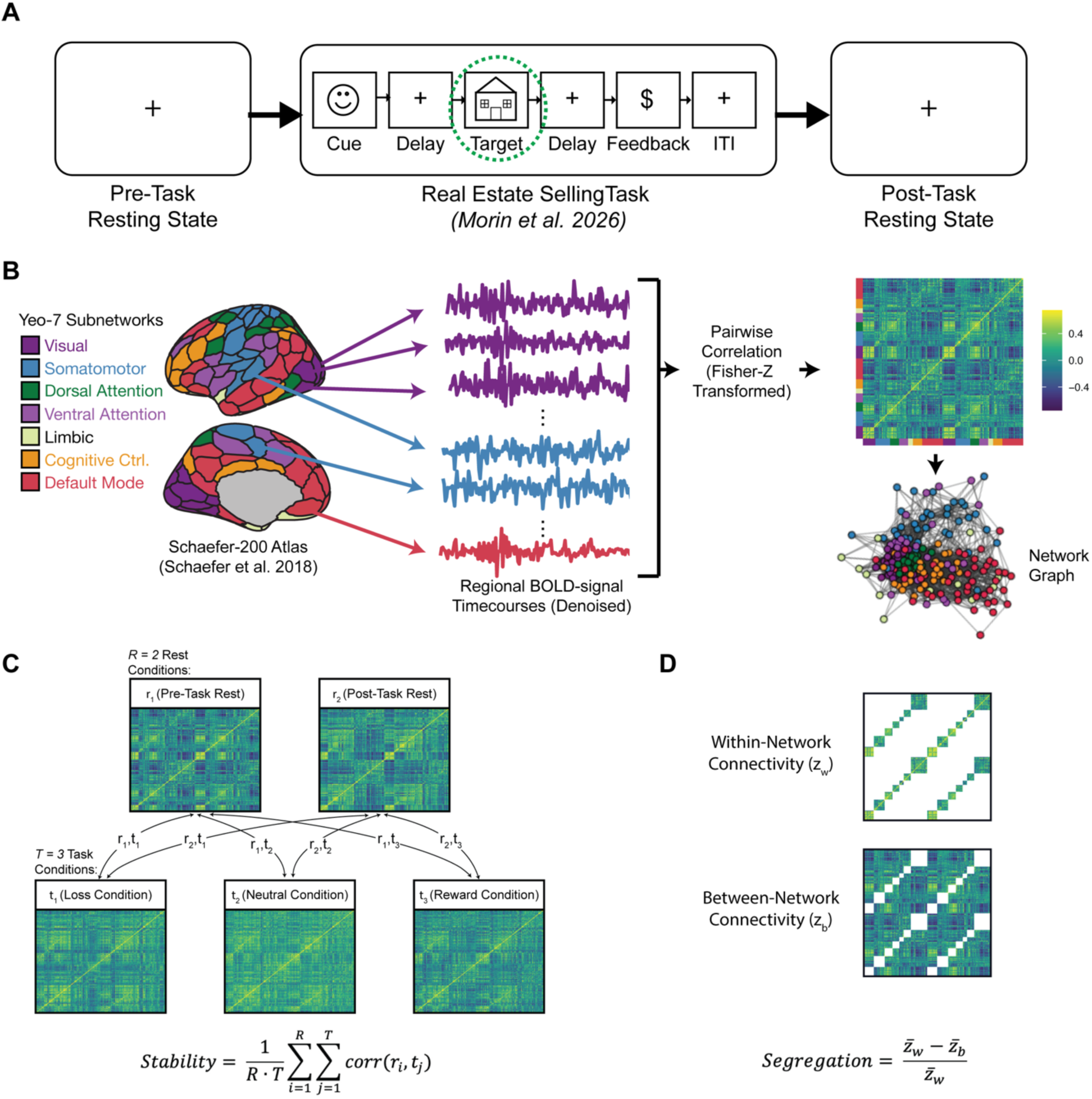
Experimental Overview. **(A)** Participants completed fMRI during resting state, the real-estate selling task, and again during a second resting state. **(B)** Functional networks were constructed for resting and task states by extracting the mean time series from each region of the Schaefer-200 cortical atlas and calculating the Fisher-Z transformed Pearson correlation between pairs of time series. For resting states, the full time series was used. For task states, condition-specific weighted timeseries were used. Denoising procedures specific to the resting and task conditions are described in the Methods. **(C)** Network stability was calculated as the mean Pearson correlation (corr(r_i_, t_j_)) between rest network matrices, r_i_, and task network matrices, t_j_. **(D)** Network segregation was calculated as the difference of between-and within-network connectivity, normalized by within-network connectivity, using the Yeo-7 network parcellations assigned to each node in the atlas.

### Older adults show reduced network stability and segregation

To quantify network stability across rest and task states, we calculated the Fisher-Z transformed Pearson correlation between vectorized functional connectivity matrices derived from resting-state and task-based fMRI (see Methods). Network segregation was calculated as the normalized difference between within-network and between-network connectivity, based on the Yeo 7 network parcellation (see Methods) (Chan et al., 2014). Preliminary analyses found no effect of specific task condition (reward, neutral, or loss) on network measures, so measures were collapsed across resting and task states (see **Supplementary Figure S1**). Although differences in network metrics were observed between pre-and post-task resting state, these differences did not appear to influence trends associated with biomarker group, dopamine synthesis capacity, or tau burden. For simplicity in the main text, we consider the average effect of resting state across the pre-and post-task rest conditions. For completeness, results separated out by pre-and post-task resting state are displayed in **Supplementary Figures S1, S2, S3, and S4**.

Differences in network metrics across biomarker groups were tested using linear models controlling for sex, years of education, and head motion [metric ∼ biomarker_group + sex + years_edu + head_motion]. Considering network stability, PiB+ older adults showed significantly lower network stability compared to young adults (b =-0.76, t =-2.53, p =.01), but PiB-older adults did not differ significantly from young adults (b =-0.50, t =-1.81, p =.08). In a post-hoc pairwise t-test the PiB+ and PiB-groups did not significantly differ in network stability (b = 0.26, t = 0.81, p =.70). Considering network segregation during rest, older adults in both the PiB+ group (b =-0.98, t =-3.76, p <.001) and the PiB-group (b =-1.27, t =-5.96, p <.001) showed lower network segregation compared to young adults. In a post-hoc pairwise t-test, the PiB+ and PiB-groups did not significantly differ in network segregation at rest (b =-0.29, t =-1.22, p =.45). Considering network segregation during the task, no significant difference was detected between young adults and the PiB-group (b =-0.54, t =-1.77, p =.08) or between young adults and the PiB+ group (b = 0.56, t =-1.77, p =.08). A post-hoc pairwise t-test also found no significant difference between the PiB+ and PiB-groups in network segregation during task (b = 0.03, t = 0.09, p = 1.0) (**Figure 2**).

**Figure 2.**
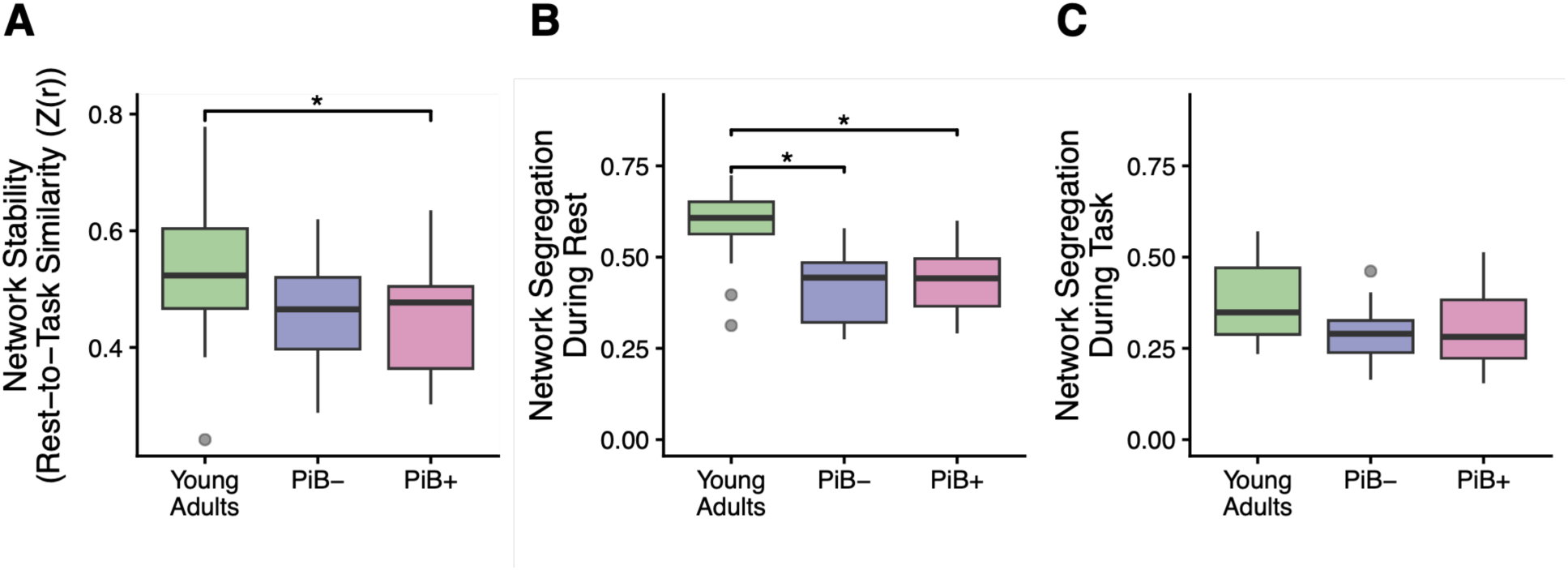
Older adults show reduced network stability and reduced network segregation. **(A)** Network stability (quantified as the Z-transformed Pearson correlation between rest and task functional connectivity matrices) was greater in young adults compared to PiB+ older adults. **(B)** Network segregation at rest was significantly higher in young adults compared to PiB-and PiB+ older adults. **(C)** Network segregation during the task showed no group differences. (* = p <.05, significant difference between biomarker groups according to a linear model controlling for sex, years of education, and head motion)

### Better Memory is Associated with Increased Network Stability and Network Segregation

Associations between network metrics (stability and segregation) and memory performance were assessed using linear models that included covariates for biomarker group, sex, years of education, and head motion [memory ∼ network metric + biomarker_group + sex + years_edu + head_motion]. The model demonstrated a significant positive association between network stability at the whole-brain level and subsequent memory scores (d’) (b = 0.34, t = 2.49, p =.02). Moreover, whole-brain network segregation during task (b = 0.34, t = 2.56, p =.01), but not at rest (b = 0.22, t = 1.12, p =.27) was significantly associated with subsequent memory (d’) (**Figure 3**). Supplementary analyses separating out the effects of pre-vs. post-task rest showed similar trends across both conditions (see **Figure S2**.)

**Figure 3.**
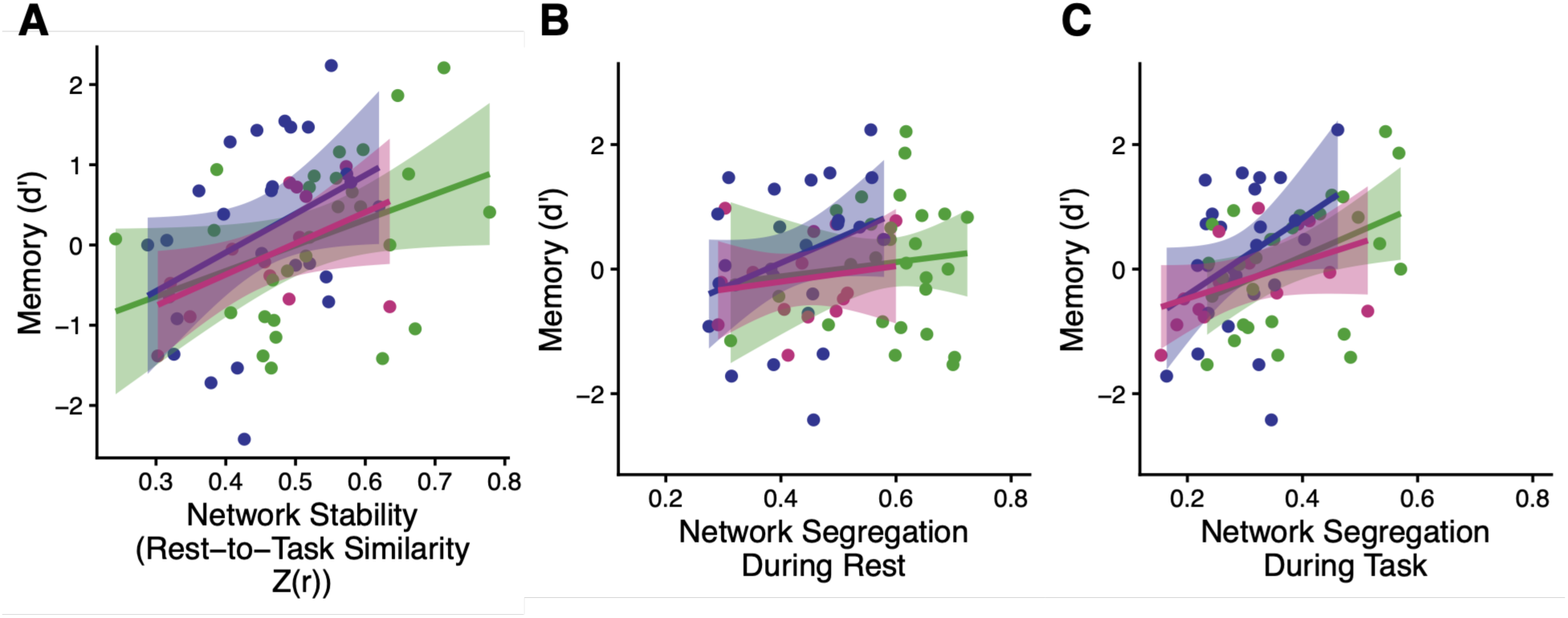
Better subsequent memory is associated with greater network stability and greater network segregation during task. **(A)** Increased network stability was associated with better subsequent memory for task stimuli (d’). **(B)** Network segregation during rest was not associated with subsequent memory scores (d’). **(C)** Increased network segregation during the task was significantly associated with better memory performance (d’). (Trend lines represent linear fit; Shaded area = 95% confidence interval; Green = Young Adults, Blue = PiB-, Pink = PiB+.)

### Interactive effect of dopamine synthesis capacity and biomarker group on network stability

Dopamine synthesis capacity was measured using [^18^F]FMT PET, with K_i_ values extracted from the ventral striatum following partial volume correction (see Methods). A linear model was used to examine interactive effects of biomarker group and dopamine synthesis capacity on network stability at the whole brain level [stability ∼ group * dopamine + sex + years_edu + head_motion]. We observed a significant interaction between dopamine synthesis capacity and biomarker group on network stability such that PiB+ older adults showed a significantly different relationship between stability and dopamine compared to younger adults (b =-0.81, t =-2.03, p =.049) (see **Figure 4**). Despite the interaction effect, which suggests that the relationship is significantly different across groups, post hoc testing with emtrends found no statistically significant relationship between dopamine synthesis capacity and whole-brain network stability within any individual group (young adults: Mean = 147.7, 95% CI = [-4.50, 39.0]; PiB-: Mean = 45.6, 95% CI = [-6.06, 27.4]; PiB+: Mean =-99.1, 95% CI = [-30.1, 5.02]).

**Figure 4.**
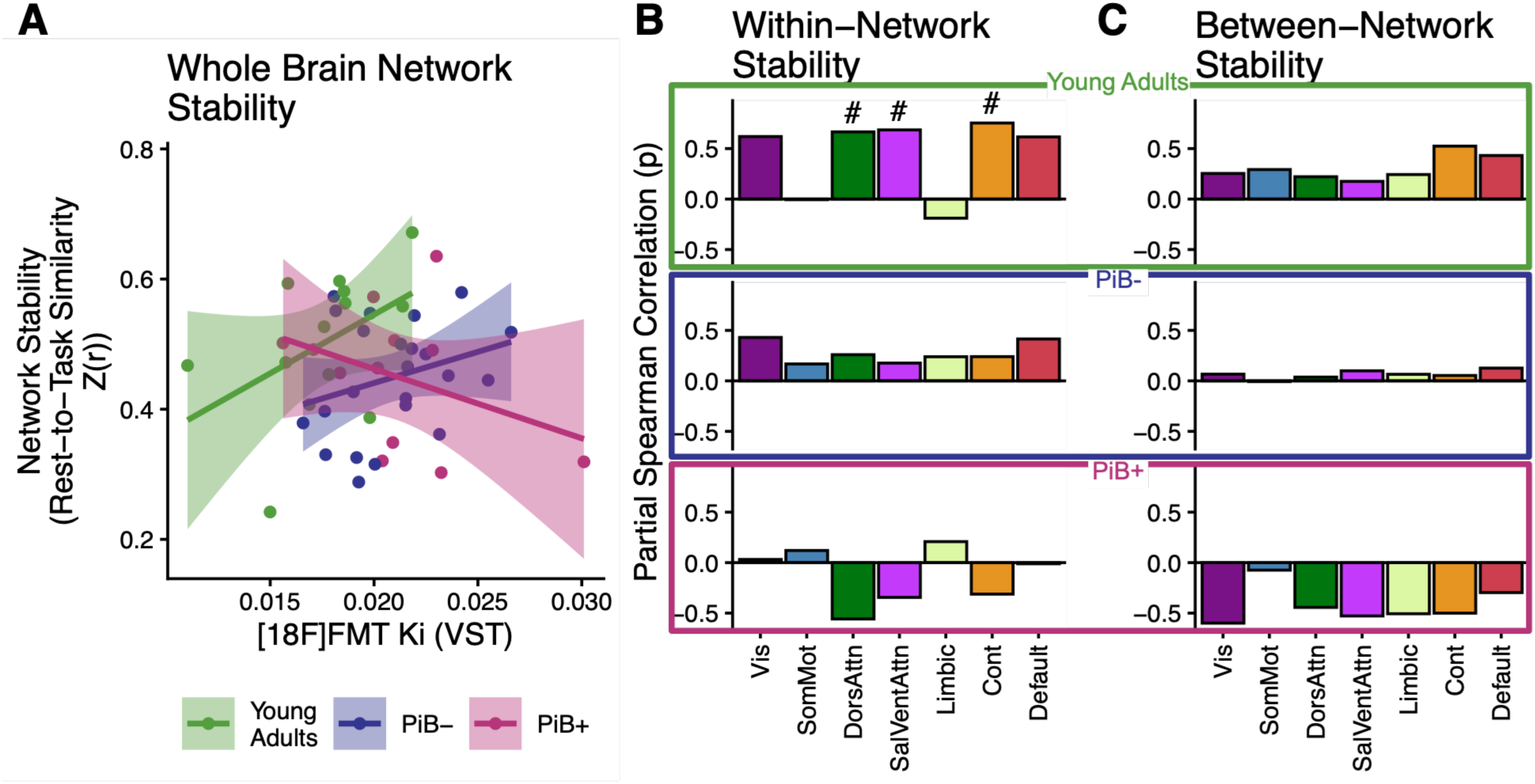
Interactive effects of dopamine synthesis capacity and biomarker group on network stability. **(A)** We observed a significant interaction between participant group and dopamine synthesis capacity on whole-brain network stability. **(B)** At the level of individual networks, young adults showed a significant positive relationship of stability within associative networks (dorsal attention, ventral attention, and cognitive control) with higher dopamine synthesis capacity in the VST, though this was not significant after correction for multiple comparisons. **(C)** Between-network connectivity was not significantly associated with dopamine synthesis capacity for any of the three subject groups. (Trend lines = linear fit, Shaded area = 95% confidence interval. Green = Young Adults, Blue = PiB-, Pink = PiB+; VST = ventral striatum, Vis = Visual, SomMot = Somatomotor, DorsAttn = Dorsal Attention, SalVentAttn = Salience/Ventral Attention, Cont = Cognitive Control; # = p <.05, uncorrected; * = q <.05, FDR corrected for multiple comparisons)

In addition to whole-brain metrics, stability and segregation can also be quantified within individual Yeo networks (e.g., visual, somatomotor) to test whether specific subnetworks show relationships between network organization and [^18^F]FMT K_i_. Subnetwork stability was calculated as rest-to-task similarity within a given network (i.e., comparing that network’s connectivity matrix at rest versus during the task). Subnetwork segregation was calculated using the connections to nodes in each subnetwork, as the difference between within-subnetwork and between-subnetwork connectivity, normalized by within-subnetwork connectivity. The partial Spearman correlation between stability and [^18^F]FMT K_i_ was calculated at the level of individual Yeo-networks for each participant group. In analyses of the young adult group, partial correlations controlled for sex, years of education, and head motion. In analyses of the older adult groups, partial correlations controlled for age, sex, years of education, and head motion. False discovery rate (FDR) correction for multiple comparisons was applied across subnetworks. We found that greater dopamine synthesis capacity in young adults was associated with increased network stability within association networks (dorsal/ventral attention and cognitive control), although these results did not survive FDR correction (visual: rho = 0.62, p =.06, q =.08; somatomotor: rho =-0.006, p =.99, q =.99; dorsal attention: rho = 0.67, p =.04, q =.08; ventral attention: rho = 0.69, p =.03, q =.08; cognitive control: rho = 0.75, p =.01, q =.08; default mode: rho = 0.62, p =.06, q =.08). In normal aging, no significant relationships between dopamine synthesis capacity and within-network stability were observed (visual: rho = 0.62, p =.06, q =.27; somatomotor: rho = 0.17, p =.50, q =.50; dorsal attention: rho = 0.27, p =.26, q =.45; ventral attention: rho = 0.18, p =.47, q =.50; limbic: rho =.24, p =.32, q =.45; cognitive control: rho = 0.25, p =.31, q =.45; default mode: rho = 0.41, p =.08, q =.27). In PiB+ older adults, we did not observe a significant relationship between dopamine synthesis capacity and within-network stability (visual: rho =-0.06, p =.89, q =.89; somatomotor: rho = 0.12, p =.78, q =.71; dorsal attention: rho =-0.58, p =.13, q =.71; ventral attention: rho =-0.42, p =.30, q =.71; limbic: rho = 0.22, p =.60, q =.84; cognitive control: rho =-0.43, p =.29, q =.71; default: rho =-0.23, p =.59, q =.84). Moreover, none of the groups showed a relationship between dopamine synthesis capacity and between-network stability. Supplementary analyses separating out the effects of pre-vs. post-task rest showed similar effects across both conditions (see **Figure S3**).

To evaluate the specificity of the dopamine-stability association, we tested whether dopamine synthesis capacity also related to rest-to-task change in network segregation (see **Figure S5**). At the whole-brain level, we did not observe an interaction of dopamine synthesis capacity and biomarker group on change in network segregation (PiB-vs. Young Adults: b =-0.25, t =-0.69, p = 0.49; PiB+ vs. Young Adults: b = 0.12, t =.33, p =.75). Exploratory analyses of subnetworks showed that in PiB-older adults, increased dopamine synthesis capacity was related to higher segregation of the dorsal attention (rho = 0.43, p = 0.04, q = 0.15) and limbic (rho = 0.61, p =.01, q =.04) networks at rest compared to the task (though the relationship with the dorsal attention network segregation did not survived corrections for multiple comparisons). No other significant associations between change in network segregation and dopamine synthesis capacity were observed.

### In PiB+ older adults, tau burden is related to task-induced collapse of network segregation

The interactive effect of dopamine synthesis capacity and biomarker group on network stability prompted us to explore whether tau burden may be associated with dysregulated network topology in the PiB+ participants. Tau pathology was quantified using [^18^F]FTP PET with standardized uptake value ratios extracted from a meta-ROI encompassing regions of the medial temporal lobe vulnerable to AD-related tau pathology (see Methods). A linear model was used to assess relationships between tau burden and change in network segregation between rest and task states at the whole-brain level [segregation_change ∼ biomarker_group * tau + age + sex + years_edu + head_motion]. We found a significant interaction between biomarker group and tau burden on change in segregation such that PiB+ older adults showed a more positive relationship between increased tau burden and change in network segregation across rest and task states (b = 0.82, t = 2.63, p =.01). Post-hoc testing with emtrends demonstrated that the association between tau burden and change in network segregation was significant in PiB+ older adults (Mean = 3.89, 95% C.I. = [1.62, 6.16]) but not in PiB-older adults (Mean =-1.53, 95% C.I. = [-5.00, 1.94]).

Exploratory follow-up analyses were conducted at the level of each Yeo-7 subnetwork using partial Spearman correlation for each group (PiB-and PiB+) controlling for age, sex, years of education, and head motion. Results were FDR-corrected for multiple comparisons across the subnetworks. We found that for PiB+ participants, tau burden was significantly associated with rest-to-task change in segregation in visual, somatomotor, dorsal attention, ventral attention, and cognitive control networks (visual: rho = 0.91, p <.001, q <.01; somatomotor: rho = 0.88, p <.01, q <.01; dorsal attention: rho = 0.74, p =.03, q =.04; ventral attention: rho = 0.81, p =.02, q =.03; limbic: rho = 0.69, p =.06, q =.07; cognitive control: rho = 0.93, p <.001, q <.01; default: rho = 0.62, p =.10, q =.10). No significant associations were observed in PiB-participants (visual: rho =-0.34, p =.14, q =.93; somatomotor: rho =-0.05, p =.84, q =.93; dorsal attention: rho = 0.02, p =.93, q =.93; ventral attention: rho =-0.11, p =.65, q =.93; limbic: rho =-0.03, p =.89, q =.94; cognitive control: rho =-0.18, p =.44, q =.93; default mode: rho =-0.17, p =.47, q =.93). Supplementary analyses separating out the effects of pre-and post-task rest showed similar effects across both conditions (see **Figure S4**).

To evaluate the specificity of the tau-segregation association, we tested whether tau burden was also related to network stability (see **Figure S6**). At the whole-brain level, no significant interactive effect of biomarker group and tau-PET measures on network stability was observed (b =-0.51, t =-1.4, p = 0.17). Exploratory analyses of subnetworks showed significant negative associations between within-network stability and tau-PET measures in the PiB-group (cognitive control: rho =-0.45, p =.047, q =.33) and PiB+ group (cognitive control: rho =-0.72, p = 0.04, q = 0.15; default mode: rho =-0.85, p <.01, q =.06), but these relationships did not survive correction for multiple comparisons. No significant associations were detected for between-network stability and tau-PET measures at the level of specific Yeo subnetworks.

### Network stability is related to network segregation

Conceptual frameworks propose that increased network segregation might predispose the brain to greater stability during transitions between rest and task (Cole et al., 2021; Hughes et al., 2025). To explore whether this framework was supported in our data, we modeled the direct relationships between network stability and each of the network segregation measures (segregation during rest, segregation during task, and change in segregation) at the whole brain level (see **Figure 6**). Inspired by prior work showing an association between network segregation at rest and network stability in young adults, but not older adults (Hughes et al., 2025), models tested for an interaction between biomarker group and network segregation metric on measures of network stability. Models controlled for biomarker group, sex, years of education, and head motion (network_stability ∼ network_segregation * biomarker_group + sex + years_edu + head_motion). We did not observe any significant interaction effects, but network segregation during rest (b = 0.56, t = 2.15, p =.04), network segregation during task (b =.60, t = 4.28, p <.001) and change in segregation (b =-.43, t =-2.45, p =.02) showed significant associations with network stability.

**Figure 5.**
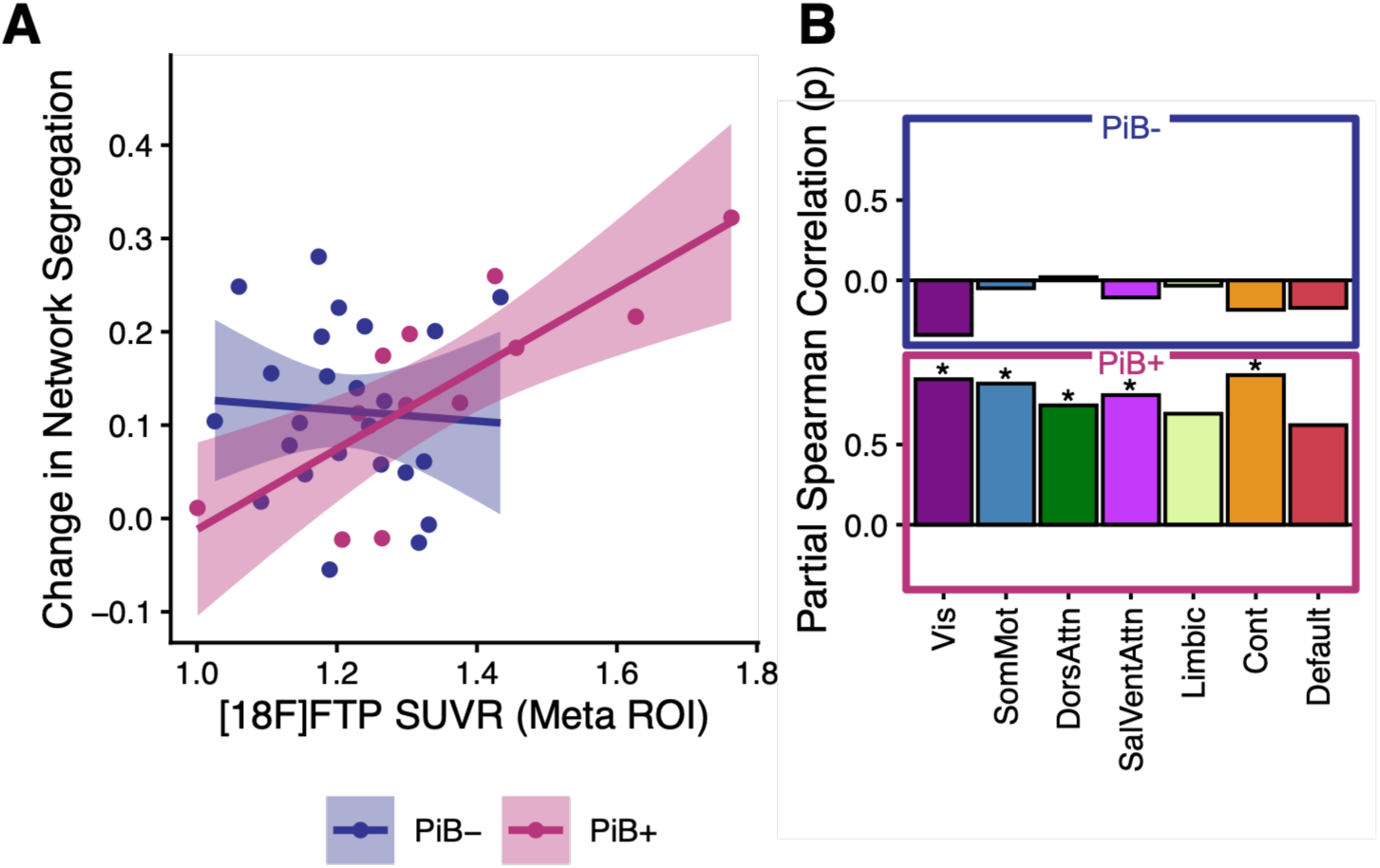
Tau burden predicts higher network segregation at rest compared to task in PiB+ older adults. **(A)** Increased tau burden was associated with a larger drop in segregation from rest to task in the PiB+ group, but not the PiB-group. **(B)** This relationship was significant across visual, somatomotor, dorsal attention, salience/ventral attention, and cognitive control Yeo-networks for the PiB+, but not the PiB-group. (Trend lines = linear fit, Shaded area = 95% confidence interval, Blue = PiB-, Pink = PiB+; Vis = Visual, SomMot = Somatomotor, DorsAttn = Dorsal Attention, SalVentAttn = Salience/Ventral Attention, Cont = Cognitive Control; # = p <.05, uncorrected; * = q <.05, FDR corrected for multiple comparisons).

**Figure 6.**
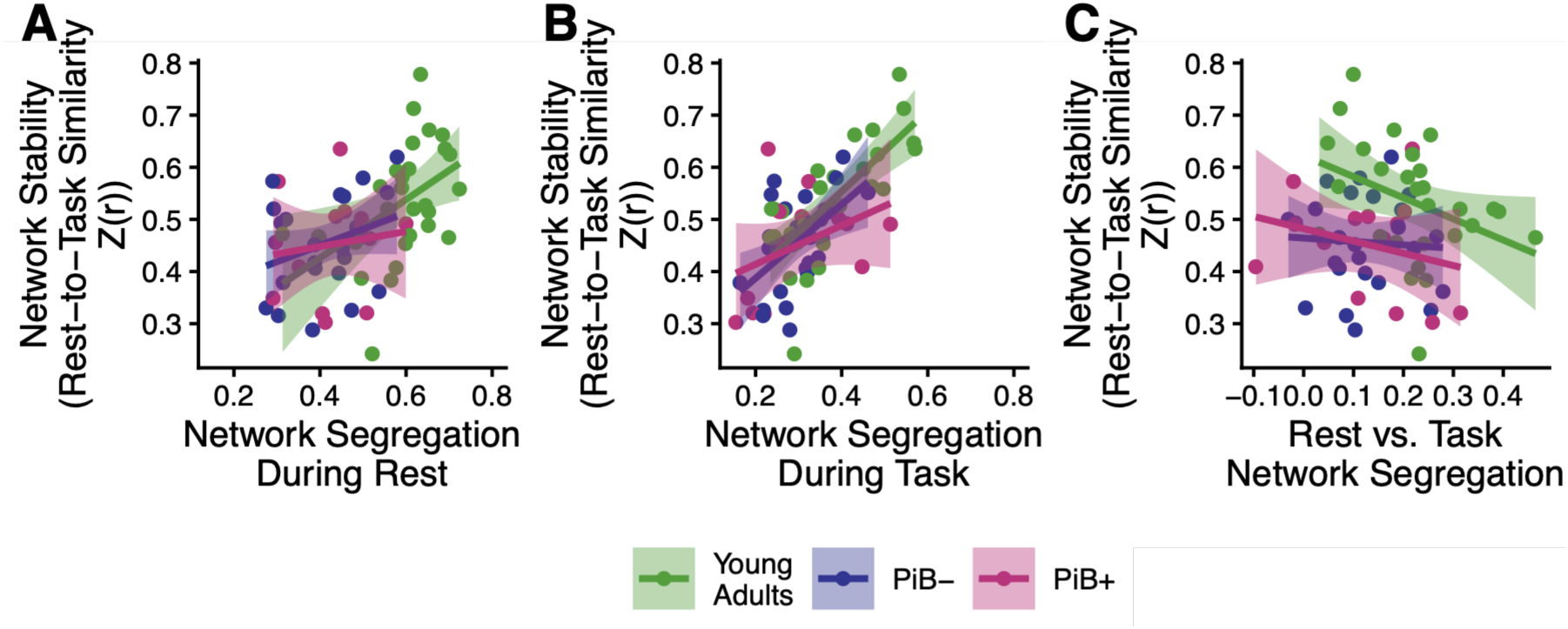
Network stability is related to network segregation. Network stability is plotted against network segregation during rest **(A)**, network segregation during task **(B)**, and change in network segregation **(C)**. (Trend lines = linear fit, Shaded area = 95% confidence interval. Green = Young Adults, Blue = PiB-, Pink = PiB+)

## Discussion

This study combined PET markers of dopamine synthesis capacity ([^18^F]FMT), tau burden ([^18^F]FTP), and amyloid-β status ([^11^C]PiB) with resting state and task-based fMRI to test how neuromodulatory and pathological processes shape large-scale network dynamics across the adult lifespan. We found that higher ventral striatal dopamine synthesis capacity was associated with greater network stability within associative networks in young adults, but that this relationship was attenuated in older adults and absent in PiB+ participants. In PiB+ older adults, we found that higher tau burden was associated with a task-induced collapse of network segregation across large-scale networks. Importantly, both greater network stability and greater task-related segregation predicted better subsequent memory, linking these network-level effects to cognition. Together, these results suggest that age-and pathology-related changes in network organization reflect opposing influences of dopamine and tau, highlighting rest-to-task network dynamics as a promising intermediate phenotype for understanding heterogeneity in cognitive aging.

### Age-related effects on rest-to-task network stability

Previous work studying young adult participants has documented the selective integration of networks elicited by cognitive tasks, and has demonstrated that improved performance on a task is related to greater network stability across resting and task states (Cohen & D’Esposito, 2016; D. H. Schultz & Cole, 2016; Thiele et al., 2022). These results have been extended to the field of cognitive aging with large-scale studies demonstrating that network segregation is reduced in older age, and that reduced segregation is associated with poorer cognitive outcomes (Betzel et al., 2014; Chan et al., 2014, 2021). Moreover, increased age is associated with reduced network stability, which is in-turn related to lower scores on cognitive measures of working memory (Yue et al., 2024), social cognition (Hughes et al., 2025), and episodic memory for objects (Cassady et al., 2023). Our results replicated findings that older adults show lower network stability and segregation compared to young adults, and that individual differences in these network metrics are related to task performance.

To measure task performance, we chose to examine subsequent memory for stimuli, given established links between AD pathology and memory in aging. Our measure of task-based functional connectivity was derived from the target-period of the task when house stimuli were incidentally encoded by participants. Although these methods maximize our ability to measure network dynamics during memory encoding, they do result in a “fragmented” time series composed of short temporal epochs from individual trials. Future studies employing a blocked design, time-windowed connectivity, and/or dynamical systems modeling may clarify whether network dynamics are entirely stable throughout different phases of the task, or whether they temporarily re-enter a rest-like state during successful encoding. Future studies should also consider matching the scanning parameters across rest and task states (e.g. voxel size, TR, slice acquisition framework) as an extra measure of control.

Importantly, our study extends prior work by showing that memory retrieval measured outside of the scanner is related to network dynamics measured during memory encoding inside the scanner, even when the memory encoding was incidental to the task the subject was performing. This may suggest that task-induced network integration and network stability provide domain general cognitive benefits, relevant not only to the task at hand (i.e. predicting property sales, in our task) but also extending to other cognitive domains that are simultaneously engaged (i.e. encoding tangential details of the trial). More broadly, previous studies that examined network dynamics across multiple tasks in the same individuals suggest that network stability may be a domain-general trait-like property related to overall measures of intelligence (Schultz & Cole, 2016; Thiele et al., 2022). Together these findings warrant further investigation into whether network dynamics represent a state-like phenomenon specific to arbitrary tasks, a trait-like domain-general phenomenon reflective of persistent individual differences, or a combination of the two. The precise mechanisms that may influence age-related declines in network segregation and stability are not fully understood, but could be attributed to accumulated life experience, reduced neuronal integrity, and as we suggest, neuromodulatory and neuropathological factors.

### Dopaminergic support of network stability

Our results confirm and extend prior computational and empirical work demonstrating that dopamine influences the stability of task-relevant network states. Interestingly, much of this theorizing comes from work in small neural circuits (computational models of only a few neurons), but may be generalized to the kind of large-scale brain networks we can measure with fMRI due to the scale-invariant properties of the brain. Neural gain models propose that dopamine stabilizes cortical attractor dynamics, increasing the signal-to-noise ratio in neurons receiving afferent input from brainstem nuclei, thereby promoting consistent patterns of functional connectivity and increased network integration during cognitive engagement (Shine et al., 2018). Previous studies testing associations with pharmacological manipulation (Garrett et al., 2015), genetic polymorphisms (Braun et al., 2021), and BOLD variability (Garrett et al., 2022) have implicated dopamine in shaping large-scale network segregation and stability, but have largely relied on indirect proxies for measuring dopamine function. By using [^18^F]FMT PET, we provide direct evidence for an interactive relationship between dopamine synthesis capacity and whole-brain network stability, such that higher ventral striatal dopamine synthesis capacity was associated with greater network stability in young adults, but that this relationship was attenuated in PiB-older adults and absent in PiB+ participants. Moreover, we showed preliminary evidence that individual differences in dopamine synthesis capacity are associated with greater stability within dorsal attention, ventral attention, and cognitive control networks in young adults (though these results were not significant after correction for multiple comparisons and should be replicated in a larger sample).

Because the task couples reward learning (monetary feedback) with subsequent recognition memory, it provides a motivation-relevant context in which dopaminergic support for network stability may be especially consequential for successful memory encoding. Notably, these associations were attenuated in PiB-and absent in PiB+ older adults, suggesting that dopaminergic support for network stability may diminish as aging-and AD-related pathology accumulate. One interpretation is that dopaminergic support for network stability is strongest in young adulthood and becomes more difficult to detect in older adulthood due to dopamine-system dysfunction and increased inter-individual heterogeneity. Another possibility is that tau-mediated dysfunction in medial temporal brain regions disrupts the ability of dopaminergic projections (originating in the ventral tegmental area and innervating the ventral striatum and hippocampus) to effectively stabilize network transitions during cognitive processing. Future studies incorporating tasks that probe executive control, non-rewarded memory, or sensory processing will be important for determining whether dopaminergic modulation of rest-to-task network stability generalizes across cognitive domains.

Although we limited this investigation to measures of dopamine synthesis capacity in the ventral striatum using [^18^F]FMT, network dynamics are likely influenced by other facets of the dopamine system (i.e. dopamine release, receptor density) in addition to other neuromodulatory systems (e.g. serotonin, norepinephrine, etc.). Future studies might employ radiotracers with uptake in cortical regions to see if local receptor density is associated with regional network dynamics. Existing work using [^11^C]NNC112 found links between D1 receptor density in lateral prefrontal cortex and connectivity strength within networks (Roffman et al., 2017). Atlas-based methods for mapping the normative spatial distribution of receptor systems may also offer insight on the relative influence of multiple neuromodulatory systems on brain network dynamics (Hansen et al., 2022).

### Tau-driven collapse of network segregation in PiB+ older adults

In contrast to the stabilizing role of dopamine, we observed that tau burden in PiB+ older adults was associated with a larger reduction in network segregation during task engagement (i.e., a greater drop in segregation between rest and task states). This finding is broadly consistent with prior work linking tau and Aβ to altered functional connectivity, particularly with medial temporal and default-mode regions (Berron et al., 2020; Cassady et al., 2021; Sepulcre et al., 2017), and with evidence that early AD pathophysiology can be accompanied by widespread hyperconnectivity (A. P. Schultz et al., 2017). Related work also suggests that impaired clearance of Aβ and tau pathology may contribute to unconstrained propagation of activity from associative to sensory regions, a feature consistent with the idea that high-tau individuals exhibit task-induced collapse of network segregation (F. Han et al., 2025).

Critically, tau related most strongly to a decrease in network segregation between rest and task states (see **Figure 5**), but showed no association with whole-brain network stability (see **Figure S6**). This pattern is plausible because these two metrics of functional network dynamics are sensitive to different features of the connectivity matrix: segregation indexes the balance of within-versus between-network coupling within a given state, whereas stability summarizes the overall similarity of whole-matrix edge patterns across states. A systematic reduction in segregation can arise from a relatively uniform increase in between-network coupling (or decrease in within-network coupling) during the task, altering modular boundaries without substantially reshaping the rank-order pattern of edges that dominates the stability correlation. In this sense, reduced segregation reflects a bias toward unconstrained integration during task engagement, not necessarily greater “flexibility” in the form of large-scale reconfiguration of connectivity patterns between rest and task. At the level of subnetworks, we observed a negative association between tau burden and stability within the default mode network in PiB+ older adults, but this did not remain significant after correction for multiple comparisons (see **Figure S6b**). This result is intriguing, as it may reflect previous findings that tau burden in the posterior memory network (which overlaps with the default mode network) is associated with reduced stability of this network (Cassady et al., 2023).

Previous work suggests that greater network segregation during rest may facilitate network stability during transitions between resting and task states (Cole et al., 2021; Hughes et al., 2025). Within this framework, a more segregated resting-state architecture may provide a task-ready organizational scaffold, such that relatively modest shifts in network configuration are required to achieve task-appropriate levels of activation in relevant regions. Consistent with this interpretation, we found significant associations between measures of network segregation and network stability in our study (**Figure 6**). We propose that in young adults and in cognitively normal aging, network segregation promotes stable rest-to-task transitions and is supported, at least in part, by the dopamine system. Upon engaging with a task, this architecture may allow task-relevant regions to selectively integrate while preserving network boundaries. However, in the context of AD-related pathology, this relationship may break down. Elevated tau burden may increase neural or network-level noise, permitting unconstrained propagation of task-evoked activity across the connectome and leading to excessive, rather than selective, network integration.

### Converging and competing influences of dopamine and tau

A key contribution of this study is demonstrating that dopaminergic function and tau pathology influence network dynamics through partially opposing mechanisms. Dopamine appears to enhance stability and preserve segregation, particularly within associative networks, whereas tau undermines segregation and disrupts task transitions. Although the richness of this multimodal imaging dataset offers a unique window into these mechanisms, the modest sample size underscores the need for replication in larger cohorts with sufficient power to characterize regional specificity and individual differences. Longitudinal multimodal imaging will be particularly important for determining whether dopaminergic decline and tau accumulation unfold in coordinated or independent trajectories, and for understanding how their interaction contributes to emerging network instability and cognitive decline. Together, these competing forces may help explain heterogeneous aging trajectories: in some older adults, preserved dopaminergic function may buffer the impact of early pathology, whereas in others, elevated tau may override neuromodulatory support, leading to network-level dysregulation.

## Methods

### Participants

Participants included fifty older adults (ages 60 – 80) and thirty young adults (ages 18 – 35) from the Berkeley Aging Cohort Study. Older adults were characterized as cognitively normal using an interview and test battery including subsets of the Wechsler Adult Intelligence Scale-Revised (Wechsler, 1987) and the Mini Mental State Exam (MMSE) (Folstein et al., 1975). Older adults scored at least 25 on the MMSE. All participants provided informed consent and study procedures were approved by the Institutional Review Boards at University of California, Berkeley and Lawrence Berkeley National Laboratory. All participants completed an fMRI scanning session, during which they completed both resting state and task-based scans (described below). Following the fMRI session, participants completed a surprise memory test for stimuli presented during the scan. Of the older adult participants, four were excluded for poor quality behavioral data (two failed to learn the task, one responded “Yes” to remembering every stimulus, and one withdrew early after only completing two runs of the task). Additionally, five older adults and two younger adults were excluded from all analyses due to excessive head motion (> 20% time frames censored during at least one condition: pre-task rest, post-task rest, reward, neutral, or loss). Many participants also completed PET imaging scans to measure dopamine synthesis capacity ([^18^F]Fluoro-L-*m*-tyrosine (FMT)) (n = 13 younger adults and 37 older adults who also had acceptable fMRI data), tau ([^18^F]Flortaucipir (FTP)) (n = 37 older adults who also had acceptable fMRI data), and Aβ ([^11^C]Pittsburgh Compound-B (PiB)) (n = 49 older adults; one participant without PiB data was assumed PiB-). Demographic information for the final sample included in each set of analyses is presented in **Table 1**.

### Experimental Procedure

Participants completed both resting state and task-based fMRI. Two runs of resting state fMRI were collected, one prior to the task, and one afterwards. Resting state scans were 5 minutes 20 seconds minutes in length. Participants were asked to lie still with their eyes open and a fixation cross was displayed. Four task-based scans, each approximately 9 minutes 50 seconds in length, were conducted during which participants completed the “Real Estate Selling Task” (Morin et al., 2026). Briefly, during the real estate selling task, participants were told they owned a portfolio of houses and worked with three real estate agents to sell them. Each agent had a distinctive outcome pattern: one typically sold for a profit (reward agent), one always broke even (neutral agent), and one typically sold at a loss (loss agent). Participants were instructed to learn and label each agent’s identity (reward/neutral/loss) based on repeated feedback. Each trial consisted of: (i) Cue phase (2.0s): a real estate agent was shown and participants identified the agent as “reward,” “neutral,” or “loss” using one of three buttons; (ii) Delay phase (2.4–4.8s); (iii) Target phase (3.0s): A house image appeared, outlined in green (profit), gray (break-even), or red (loss) to indicate the outcome of the sale; (iv) Feedback phase (1.0s): Monetary outcome displayed: +$1 (reward),-$0.50 (loss), or “=” (neutral); and (v) Inter-trial interval (2–10s, jittered). Trial outcomes were predetermined to optimize orthogonality across conditions for event-related fMRI analyses. If a participant failed to respond during the cue phase, then the target phase showed a house overlaid with “?” and a blank screen was displayed during the feedback phase. These non-response trials were marked as incorrect. Agent-outcome contingencies were: (i) Reward agent: 80% profit, 20% loss; (ii) Loss agent: 80% loss, 20% profit; (iii) Neutral agent: 100% break-even. Participants were not informed of these probabilities and were told to categorize agents based on their typical outcomes, not to predict individual trials.

### Memory Scores

Memory for the house images was assessed approximately 30 minutes after MRI scanning. Participants saw 72 house images and indicated whether they had previously seen each image: “Yes,” “Likely Yes,” “Likely No,” or “No.” Of the stimuli shown, 16 were lure images of houses that they had never seen before, and 56 were images of the houses previously displayed during the probabilistic learning phase (16 reward, 16 neutral, 16 loss, and 8 “surprise trials” that were unexpectedly sold at a profit/loss by the reward/loss realtor). Surprise trials were omitted from all memory analyses. Due to known effects of age on false-memory recall, we employed d’ as a measure of memory ability to account for both correct recognitions (hits) and false alarms, rather than using simple memory-accuracy measures (for review, see Devitt & Schacter, 2016). Analyses of d’ focused on high-confidence memory trials rated ‘Yes/No’. Memory performance was scored using d’, calculated as the difference between the normalized hit rate and the normalized false alarm rate:

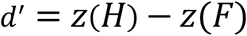

where *H* is the proportion of previously-seen house images that a subject correctly identified as remembered with high confidence (‘Yes’), and *F* is the proportion of lure house images that a subject mistakenly identified as remembered with high confidence (‘Yes’).

### FMT PET Acquisition, Reconstruction, and Processing

The irreversible PET tracer [^18^F]Fluoro-l-m-tyrosine (FMT) was used to measure dopamine synthesis capacity. [^18^F]FMT was synthesized at Lawrence Berkeley National Laboratory, as previously described (VanBrocklin et al., 2004). Participants were injected with ∼2.5 mCi of [^18^F]FMT as a bolus in an antecubital vein and scanned using a Biograph Truepoint 6 PET/CT (Siemens Medical Systems). Participants took oral carbidopa (2.5 mg/kg) approximately one hour before scanning to limit the peripheral metabolism of [^18^F]FMT. Dynamic acquisition frames were collected over a 90-minute period in 3D mode (25 frames total: 5 × 1 min, 3 × 2 min, 3 × 3 min, 14 × 5 min). Reconstruction of the scans involved an ordered subset expectation maximization algorithm with weighted attenuation, correction for scatter, and smoothing with a 4 mm full-width-at-half-maximum (FWHM) kernel. [^18^F]FMT data were processed using SPM12 software (http://www.fil.ion.ucl.ac.uk/spm). Images were realigned to the middle (12^th^) frame to correct for motion between frames, and the first 5 images were summed before realignment. PET images were coregistered to the T_1_-weighted structural scan using the mean of the frames corresponding to the first 20 min of data acquisition as a target. Patlak plotting was used to perform graphical analysis for irreversible tracer binding (Patlak & Blasberg, 1985) with posterior gray cerebellum as the reference region. K_i_ maps were generated from PET frames corresponding to 25 min and 90 min (Ciampa et al., 2022; Parent et al., 2022), which represent the amount of tracer accumulated in the brain relative to the reference region. K_i_ can be expressed as *K*_i_ = *k*_2_*k*_3_/(*k*_2_ + *k*_3_), where *k*_2_ is the rate constant for the return of free [^18^F]FMT from brain back to plasma and *k*_3_ is the rate constant for the trapping of brain [^18^F]FMT by aromatic amino acid decarboxylase. These images are comparable to *K*_i_ images obtained using a blood input function but are scaled to the volume of tracer distribution in the reference region; we use K_i_ here for simplicity. Partial volume correction (PVC) was applied to limit contributions of surrounding gray matter, white matter, and CSF to the measurement of striatal [^18^F]FMT K_i_. PVC was applied to K_i_ images using the Geometric Transfer Matrix approach (Rousset et al., 1998). Striatal regions of interest (ROIs) were manually drawn on each participant’s T_1_-weighted structural scan as previously described (Mawlawi et al., 2001) using Mango software. To apply the PVC in native space, we used FreeSurfer-generated ROIs for gray matter cortical and subcortical regions, white matter, and CSF with manually drawn striatal ROIs substituting for the automated striatal segmentation. The striatal region of interest for this study was the ventral striatum (VST).

### FTP & PiB PET Acquisition, Reconstruction, and Processing

β-amyloid (Aβ) status was determined for a subset of older adult participants using [^11^C]Pittsburgh compound B ([^11^C]PiB) PET. Tau burden was also determined for a subset of older adult participants using [^18^F]Flortaucipir ([^18^F]FTP). All [^11^C]PiB and [^18^F]FTP scans occurred within three years of a subject’s fMRI scan. A detailed description of PiB and FTP acquisition has been published previously (Schöll et al., 2016). After PiB injection, dynamic frames were collected for 90 min. These frames were realigned, co-registered, and resliced to the 3T structural T1. Distribution volume ratio (DVR) was calculated using a Logan graphical analysis (Logan, J et al., 1990) on frames corresponding to 35–90 min post-injection using a cerebellar gray reference region. PiB status was determined using a positivity threshold of 1.065 on mean gray matter DVR (Villeneuve et al., 2015). [^18^F]FTP Standardized uptake volume ratio (SUVR) images were created based on mean tracer uptake over 80–100 min post-injection, normalized using an inferior cerebellar gray reference region. As previously described, SUVR images were partial volume corrected on native space FreeSurfer-derived ROIs (Baker et al., 2017). Tau-PET analyses focused on a voxel-weighted meta-ROI consisting of entorhinal, amygdala, parahippocampal, fusiform, inferior temporal, and middle temporal regions, previously defined in (Ciampa et al., 2022).

### MRI Acquisition

Structural and functional MRI data were acquired on a 3-Tesla Siemens TIM/Trio scanner located at the Henry H. Wheeler Jr. Brain Imaging Center in Berkeley, California. A 32-channel head coil was used for all scans. Whole-brain high resolution T1-weighted volumetric magnetization prepared rapid gradient echo image (MPRAGE) scans were acquired for each subject (voxel size = 1mm isotropic, TR = 2300ms, TE = 2.98ms, matrix = 256 x 240, sagittal plane, 160 slices). For resting state and task-based fMRI scans, T2*-weighted echo-planar images were acquired using a multiband sequence (slice acceleration factor, 4) [Resting State: TR = 1067ms, TE = 31.2ms, Flip Angle = 45°, partial-Fourier acquisition; Task-Based: TR = 2400ms, TE= 36ms, Flip Angle = 45°, partial-Fourier acquisition]. For resting state fMRI, a total of 60 slices were acquired in sequential order, covering the whole brain. 300 volumes were acquired with 2.6mm isotropic voxels (matrix size 80 x 80) and the z-axis was aligned to the AC-PC line. For task-based fMRI, a total of 88 slices were acquired in an interleaved order with a step-size of 9, covering the whole brain. 246 volumes were acquired with 1.54mm isotropic voxels (matrix size 138 x 138) and the z-axis was aligned to the AC-PC line. Two opposite phase-encoded EPI fieldmaps (one anterior-to-posterior, the other posterior-to-anterior) were also collected for each subject to be used for distortion correction of the functional images [TR = 16,000ms, TE = 111ms, flip angle = 90°].

### fMRI Preprocessing

MRI preprocessing was performed using fMRIPrep (22.0.2) which is based on NiPype (1.8.5) (Esteban et al., 2019). Preprocessing steps included skull stripping, distortion correction using the phase-encoded EPI fieldmaps, co-registration of each subject’s functional data with their anatomical scan using boundary-based registration with six degrees of freedom, estimation of head-motion parameters, slice-time correction, and spatial normalization. Full details of the reproducible fMRIPrep pipeline are described in the **Supplementary Methods**.

### Functional Connectivity

Following minimal preprocessing with fMRIPrep, resting state and task scans were denoised using AFNI (Cox, 1996). For all rest and task fMRI scans, confound regressors included six head motion parameters (x, y, and z translation and rotation), and six nuisance regressors derived from principal components analysis on BOLD signals within anatomical masks of cerebrospinal fluid and white matter (aCompCor implemented in fMRIPrep). Bandpass filtering [0.1, 0.01Hz] was performed simultaneously with the denoising regression model for both rest and task scans. Motion scrubbing was also performed during the denoising procedure for both rest and task scans, removing time points that exhibited greater than 0.5mm framewise displacement and/or greater than 5mm DVARS (spatial standard deviation of successive difference images). For the task scans, task-related signal from each of the task conditions was also removed from the time series during denoising using HRF-convolved boxcar functions. Regression of task-related signal is essential to ensure that functional connectivity observations were due to low-frequency coupling, rather than interregional similarities in task-based activation (Gonzalez-Castillo & Bandettini, 2018; Tomasi & Volkow, 2019).

Following denoising, mean time series were extracted from each region of the Schaefer-200 atlas (Schaefer et al., 2018). Prior work has demonstrated that functional connectivity metrics such as network segregation can be influenced by the length of fMRI acquisition (L. Han et al., 2024). To minimize error due to differences in frame quantity, we identified the subject with the most censored data due to head motion, and matched the length of each other subject’s data at each condition of interest. This limited the quantity of the data we analyzed to approximately 281s of each resting state run and task condition. For each of the resting state runs, 265 low-motion time frames (totaling 282s) were chosen from each ROI time series. For the task runs, 117 low-motion time frames (totaling 280.8s) were chosen from the target-period of each of the three task conditions, for each ROI time series. This resulted in 282s of pre-task rest, 282s of post-task rest, 280.8s of reward data, 280.8s of loss data, and 280.8s of neutral data.

For resting state scans, ROI-to-ROI connectivity matrices were generated by calculating the Fisher Z-transformation of the Pearson correlation coefficient for each pair of ROI timeseries. This was completed separately for the pre-task resting state and post-task resting state scans. Low motion timepoints for 282s of the resting state were selected randomly using python’s numpy.rand function to choose timepoints that were not censored. For task scans, weighted ROI-to-ROI connectivity matrices were generated for each task condition (reward, neutral, and loss). The connectivity matrices were computed using a weighted least squares linear model with weights defined by convolving a boxcar function for the condition of interest (e.g. 1 during the target period for reward trials, 0 otherwise) with a standard hemodynamic response function (Nieto-Castanon, 2020). Low-motion time points for 280.8s of each task condition were selected using python’s numpy.rand function to select timepoints that were not censored and were weighted positively by the convolved condition boxcar function and HRF.

### Calculating Rest-to-Task Network Stability

Methods for calculating functional network stability were modeled after previous work (Yue et al., 2024). For each subject and pair of conditions, network stability scores were generated by vectorizing the upper triangle of each functional connectivity matrix and calculating the Pearson correlation between the two vectors. Network stability for a single participant was defined as the mean Pearson correlation between *R* = 2 rest condition vectors (pre-task rest and post-task rest) and *T* = 3 task condition vectors (reward, neutral, and loss):

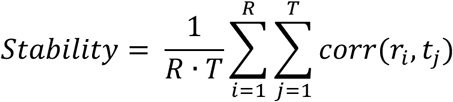

where *corr*(*r_i_*, *t*_j_) is the Pearson correlation between the functional connectivity vectors for rest condition *r_i_* and task condition *t*_j_. This was repeated at the whole brain level (whole brain network stability), for connections within each of the seven Yeo-networks (within-network stability), and for connections between each pair of Yeo-networks (between-network stability).

### Calculating Whole-Brain Network Segregation

Whole brain network segregation was calculated according to methods outlined in (Chan et al., 2014). For a specific subject and condition, segregation was defined as:

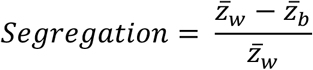

where *Z̄*_w_is the mean of the functional connectivity values connecting nodes from the same network, and *Z̄_b_* is the mean of the functional connectivity values connecting nodes from different networks. Segregation during rest was defined as the mean of the segregation measures across the two rest conditions. Segregation during task was defined as the mean of the segregation measures across the three task conditions. Change in network segregation was defined as the difference between the mean segregation of the resting state conditions and the mean segregation of the task conditions:

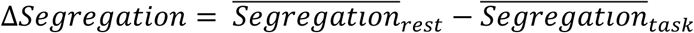

### Statistics

Linear models were used to test for relationships between network measures (stability, segregation at rest, segregation during task, and change in segregation), memory ability (d’), neurochemical markers (dopamine synthesis capacity and tau burden), and biomarker group. Each participant was assigned to one of three biomarker groups: young adults (age < 35), PiB-older adults (age > 60 with PiB mean gray matter DVR < 1.065), and PiB+ older adults (age > 60 with [^11^C]PiB mean gray matter DVR ≥1.065. Biomarker groups were coded numerically, with young adults serving as the baseline group (0). To evaluate group differences between PiB+ and PiB-older adults, post-hoc pairwise t-tests were implemented using the emmeans R package. Covariates of non-interest used in all models included sex, years of education, and head motion. Sex was coded numerically, with the female group serving as baseline (0). Because age covaried with biomarker group assignment, it was only used as a covariate in models testing for relationships within the older adult cohort. Head motion was defined as mean framewise displacement across frames included in the fMRI analyses after motion censoring and ensuring equal quantities of data across conditions and participants. For models assessing stability or change in segregation, the difference in mean framewise displacement at rest and mean framewise displacement across task conditions was used. For models assessing segregation at rest or segregation during the task, the mean framewise displacement for the corresponding rest or task condition was used. Models were fit using the lme4 package in R (version 4.0.0). Standardized beta weights for the fixed effects of interest are reported, along with T-statistics and p-values calculated using Satterthwaite’s approximation for the denominator degrees of freedom, implemented in the lmerTest R package. Code used to generate plots and run statistical analyses is available at https://github.com/tmorin01/BACS_network_dynamics

The first two sets of linear models used data from all participants with acceptable quality behavioral and fMRI data. These models assessed (1) whether network metrics differed across groups (Figure 2), and (2) whether network metrics were associated with memory ability (Figure 3).

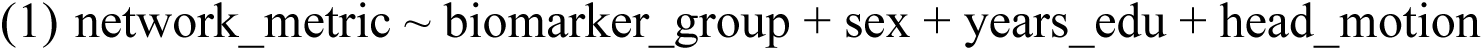

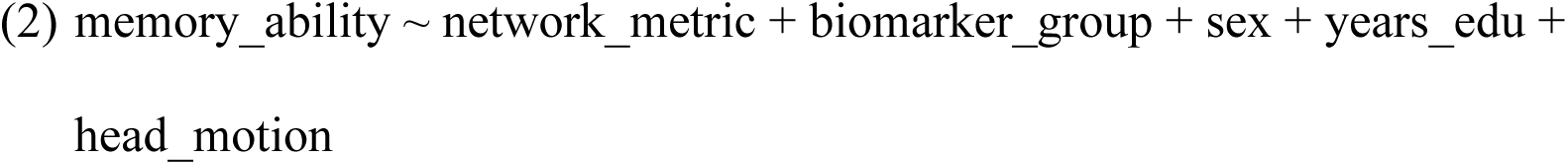

A third set of linear models used data from all participants with acceptable quality behavioral, fMRI, and [^18^F]FMT PET data. These models assessed whether (3) network metrics were associated with an interaction between biomarker group and dopamine synthesis capacity (see Figure 4 and Figure S1).

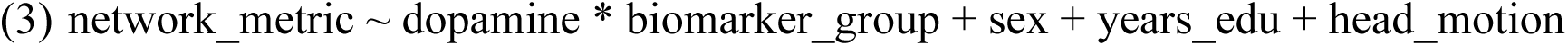

A fourth set of linear models used data from all participants with acceptable quality behavioral, fMRI, and [^18^F]FTP PET data. These models assessed whether (4) network metrics were associated with an interaction between biomarker group and tau burden (see Figure 5 and Figures S2). Note: these models included age as a covariate of non-interest because only older adult participants were included in the model. (Young adult participants did not complete FTP PET scanning.)

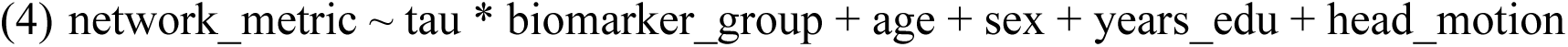

A final set of linear models used data from all participants with acceptable quality behavioral and fMRI data. These models assessed relationships between network stability and metrics of network segregation (at rest, during task, and change in segregation) (see Figure 6).

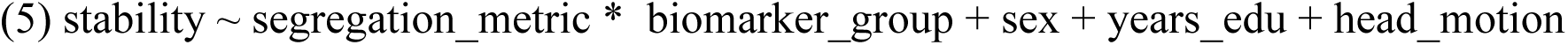

As an exploratory follow-up to model groups 3 and 4, we repeated the analysis at the level of individual Yeo-7 networks. Instead of a linear model, Spearman’s partial correlations were calculated so as not to assume linear relationships. Covariates for sex, years of education, and head motion were partialed out. Age was also partialed out for analyses of the PiB+ and PiB-groups. When examining relationships at the network level, results are presented both at a liberal threshold of p < 0.05 and after FDR-correction for multiple comparisons (q < 0.05). (Note: this procedure is modeled after Yue et al. 2024).

## Acknowledgements

This work was supported by the following grants: National Institutes of Health grants AG058748 (ASB), AG074330 (ASB), AG034570 (WJJ), AG062542 (WJJ), AG084259 (TMM), AG089518 (TMM), and Alzheimer’s Association Award AARF17-530186 (ASB). 3T MRI data were collected at the Henry H. Wheeler Jr Brain Imaging Center, which receives support from the National Science Foundation through their Major Research Instrumentation Program, award number BCS-0821855. Avid radiopharmaceuticals enabled the use of the Flortaucipir radiotracer but did not provide direct funding and were not involved in data analysis or interpretation.

Dr. Morin has served as a consultant for Rocket Science Health. Dr. Jagust has served as a consultant to Biogen, Bioclinica, Roche/Genentech, Eli Lilly, Grifols, and holds an equity interest in OptoCeutics.

