## Supplementary Materials for "Dopamine Synthesis Capacity and Tau Pathology Differentially Modulate Rest-to-Task Functional Network Dynamics"

**Supplementary Methods: fMRI Preprocessing with fMRIPrep**

MRI preprocessing was performed using fMRIPrep (22.0.2) which is based on NiPype (1.8.5) (Esteban et al., 2019). Preprocessing steps included skull stripping, distortion correction using the phase-encoded EPI fieldmaps, co-registration of each subject's functional data with their anatomical scan using boundary-based registration with six degrees of freedom, estimation of head-motion parameters, slice-time correction, and spatial normalization of the subject's anatomical and functional data to standard template space (MNI152NLin2009cAsym template from templateflow) (Ciric et al., 2021). Full details of the fMRIPrep pipeline are described below. After minimal processing with fMRIprep, we performed spatial smoothing (3mm FWHM gaussian kernel), EPI masking, and scaling so that each voxel time series had a mean of 100 using AFNI (Cox, 1996).

The following boilerplate text was produced by the fMRIPrep pipeline that was used to preprocess the MRI data in this study:

Results included in this manuscript come from preprocessing performed using fMRIPrep (22.0.2) which is based on Nipype (1.8.5).

*Preprocessing of $B_0$ Inhomogeneity Mappings:* A $B_0$ nonuniformity map (fieldmap) was estimated based on two echo-planar imaging (EPI) references with 'topup '(FSL 6.0.5.1).

*Anatomical Data Preprocessing:* The T1-weighted (T1w) image for each subject was corrected for non-uniformity (INU) with 'N4BiasFieldCorrection', distributed with ANTs 2.3.3, and used as a T1w-reference throughout the workflow. The T1w-reference was then skull-stripped with a Nipype implementation of the antsBraiNExtraction.sh workflow (from ANTs), using OASIS30ANTs as a target template. Brain tissue segmentation of cerebrospinal fluid (CSF), white-matter (WM), and gray-matter (GM) was performed on the brain-extracted T1w using 'fast' (FSL 6.0.5.1). Brain surfaces were reconstructed using 'recon-all' (FreeSurfer 7.2.0), and the brain mask estimated previously was refined with a custom variation of the method to reconcile ANTs-derived and FreeSurfer-derived segmentations of the cortical gray matter of Mindboggle. Volume-based spatial normalization to one standard space (MNI152NLin2009cAsym) was performed through using brain-extracted versions of both the T1w reference and the T1w template. The following template was used for spatial normalization: ICBM 152 Nonlinear Asymmetrical template version 2009c. (RRID: SCR_008796; TemplateFlow ID: MNI152Nlin2009cAsym)

*Functional Data Preprocessing*: For each of the four BOLD runs per subject (acquired during the Real Estate Selling Task), the following preprocessing was performed. First, a reference volume and its skull-stripped version were generated by aligning and averaging one single-band reference image (SBRef). Head-motion parameters with respect to the BOLD reference (transformation matrices, and six corresponding rotation and translation parameters) were estimated before any spatiotemporal filtering using 'mcflirt' (FSL 6.0.5.1). BOLD runs were slice-time corrected to 1.15s (0.5 of slice acquisition range 0s-2.3s) using '3dTshift' from AFNI. Slice onset times were determined from the "SliceTiming" field in the JSON sidecar file that was paired with each BOLD run. The BOLD time-series (including slice-time correction) were resampled onto their original, native space by applying the transforms to correct for head-motion. These resampled BOLD time-series will be referred to as 'preprocessed BOLD in original space' or just 'preprocessed BOLD.' The BOLD reference was then co-registered to the T1w reference using bbregister (FreeSurfer) which implements boundary-based registration. Co-registration was configured with six degrees of freedom. The BOLD-timeseries were resampled into standard space, generating a preprocessed BOLD run in MNI152NLin2009cAsym space. All resamplings can be performed with a single interpolation step by composing all the pertinent transformations (i.e. head-motion transform matrices, susceptibility distortion correction, and co-registrations to anatomical output spaces). Gridded (volumetric) resamplings were performed using ‘antsApplyTransforms’ (ANTs), configured with Lanczos interpolation to minimize the smoothing effects of other kernels. Many internal operations of fMRIPrep use Nilearn (0.9.1) mostly within the functional processing workflow. For more details of the pipeline see the section corresponding to workflows in fMRIPrep’s documentation: https://fmriprep.readthedocs.io/en/latest/workflows.html.

### Supplementary Results

#### Figure S1.

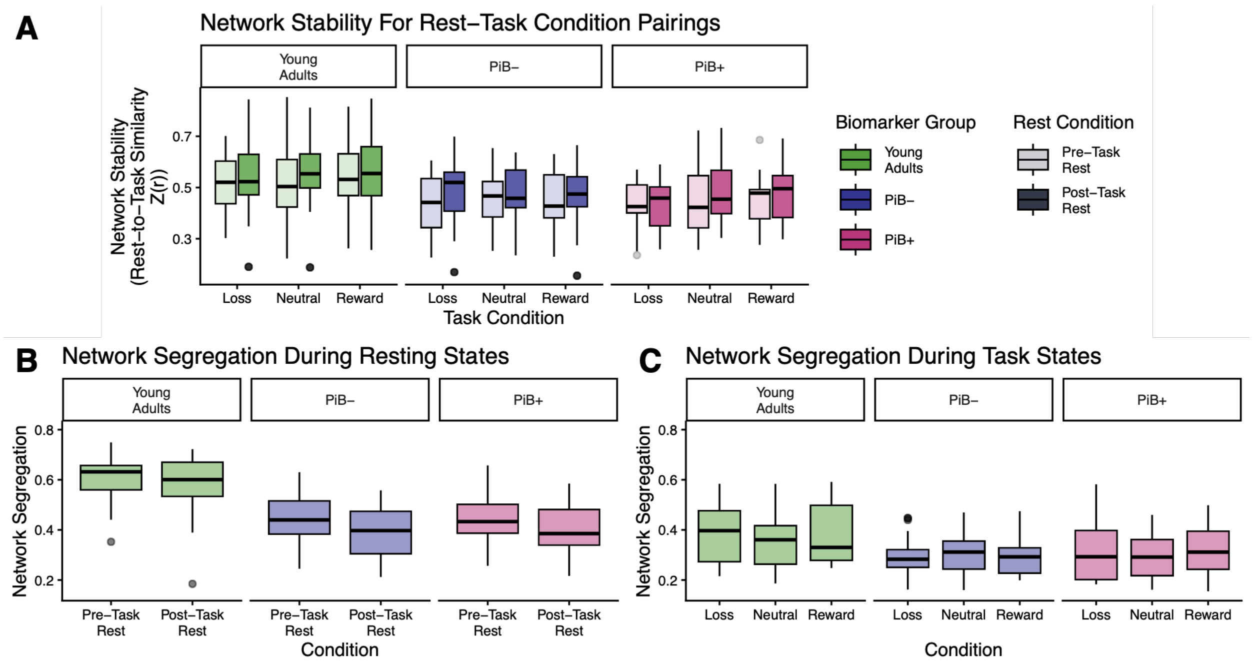


**Figure S1. Network Metrics Calculated Separately for Each Condition**. **(A)** Network stability was calculated separately for each pair of rest (pre- and post-task) and task (reward, neutral, and loss) conditions. A linear mixed effects model testing for fixed effects of biomarker group, rest condition, task condition, and controlling for sex, years of education, and head motion, with a random intercept per subject, found significant effects of biomarker group (PiB- vs. Young Adults: b = -0.56, t = -2.44, p = .02; PiB+ vs. Young Adults: b = -0.63, t = -2.41, p = .02) and rest condition (Post-Task vs. Pre-Task: b = 0.22, t = 3.81, p < .001) on network stability. There was no effect of task condition (Neutral vs. Loss: b = 0.07, t = 1.07, p = .29; Reward vs. Loss: b = 0.07, t = 0.98, p = .33) on network stability. **(B)** Network segregation during rest was calculated separately for each resting state condition. A linear mixed effects model testing for fixed effects of biomarker group and condition, and controlling for sex, years of education, and head motion, with a random intercept per subject, found significant effects of both biomarker group (PiB- vs. Young Adults: b = -1.18, t = -6.14, p < .001; PiB+ vs. Young Adults: b = -0.91, t = -3.98, p < .001), and rest condition (Post-Task vs. Pre-Task: b = -0.25, t = -2.77, p = .01) on network segregation during rest. **(C)** Network segregation during the task was calculated separately for each condition. A linear mixed effects model testing for fixed effects of biomarker group and task condition, and controlling for sex, years of education, and head motion, with a random intercept per subject, found a significant effect of biomarker group (PiB- vs. Young Adults: b = -0.63, t = -2.36, p = .02; PiB+ vs. Young Adults: b = -0.60, t = -2.04, p = .045) on segregation during task. There was no effect of task condition on network segregation (Neutral vs. Loss: b = -0.04, t = -0.49, p = .63; Reward vs. Loss: b = -0.01, t = -0.15, p = .88). These findings led us to collapse network metrics across all three task conditions. For simplicity, we also collapse across the two resting state conditions in the main text, and present the results of each rest-condition separately in the supplementary materials.

**Figure S2.**

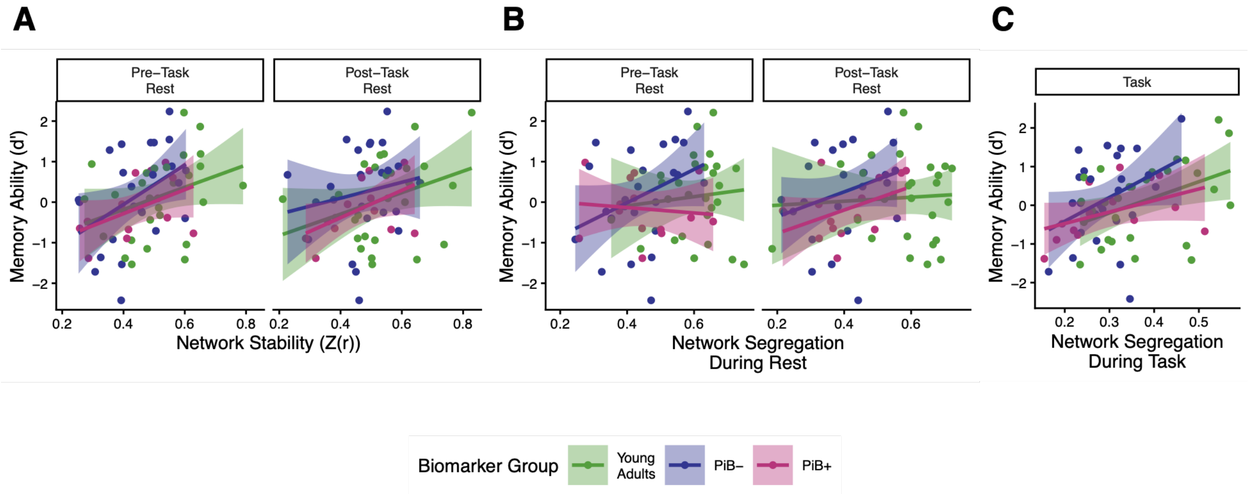


**Figure S2. Associations between memory and network metrics, separated by rest condition. (A)** Relationships between network stability and memory are plotted separately for each rest condition. A linear model testing for effects of network stability and biomarker group on memory, and controlling for sex, years of education, and head motion, found a significant effect of network stability on memory when stability was calculated for the pre-task resting state condition ($b = 0.32$, $t = 2.31$, $p = .02$). A separate model did not find a significant effect of network stability on memory when stability was calculated for the post-task resting state condition ($b = 0.27$, $t = 1.94$, $p = .06$). **(B)** Relationships between segregation at rest and memory are plotted separately for each resting state condition. A linear model testing for effects of segregation and biomarker group, and controlling for sex, years of education, and head motion, found no significant effect of segregation during pre-task rest on memory ($b = 0.19$, $t = 1.09$, $p = .28$). A separate model also did not find a significant effect of segregation during post-task rest on memory ($b = 0.17$, $t = 0.84$, $p = .41$). **(C)** The relationship between segregation during the task and memory is plotted. A linear model testing for effects of network segregation and biomarker group on memory, and controlling for sex, years of education, and head motion, found a significant effect of segregation during the task on memory ability ($b = 0.34$, $t = 2.56$, $p = .01$).

**Figure S3.**

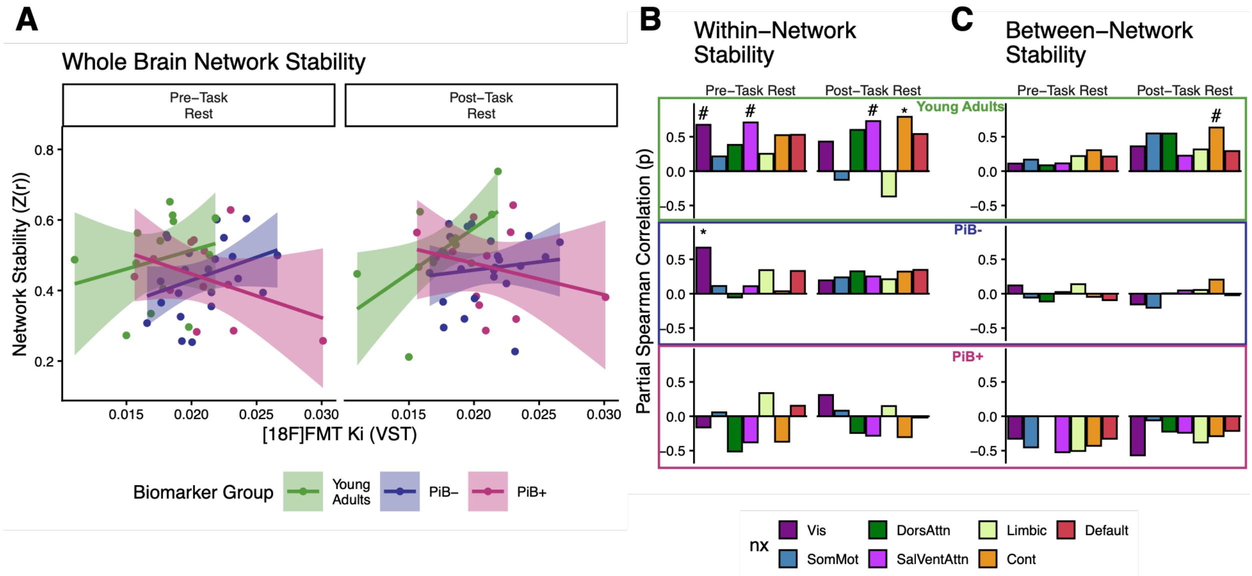


**Figure S3. Interactive effects of dopamine synthesis capacity and biomarker groups on network stability, separated out by resting state condition. (A)** Relationships between dopamine synthesis capacity in the ventral striatum ([$^{18}$F]FMT $K_i$ (VST)) and network stability are plotted for network stability calculated with pre-task rest (left) and post-task rest (right). A linear model tested for interactive relationships between dopamine synthesis capacity and biomarker group on network stability, controlling for sex, years of education, and head motion. The model found a significant interaction for network stability calculated with post-task rest, such that the PiB+ group showed a significantly more negative relationship compared to the Young Adult group ($b = -0.99$, $t = -2.54$, $p = .02$). A separate model did not find a significant interaction when network stability was calculated with pre-task rest ($b = -0.50$, $t = -1.14$, $p = .26$). **(B)** Partial Spearman correlations of within-network stability (calculated separately with each resting state condition) and dopamine synthesis capacity are plotted separately for each Yeo-7 network. Sex, years of education, and head motion were all partialed out. For the PiB- and PiB+ groups, age was also partialed out. **(C)** Partial Spearman correlation of between-network stability (calculated separately with each resting state condition) and dopamine synthesis capacity are plotted separately for each Yeo-7 condition. Sex, years of education, and head motion were all partialed out. For the PiB- and PiB+ groups, age was also partialed out. (# = $p < .05$ prior to FDR-correction; * = $q < .05$ after FDR-correction for multiple comparisons.)

**Figure S4.**

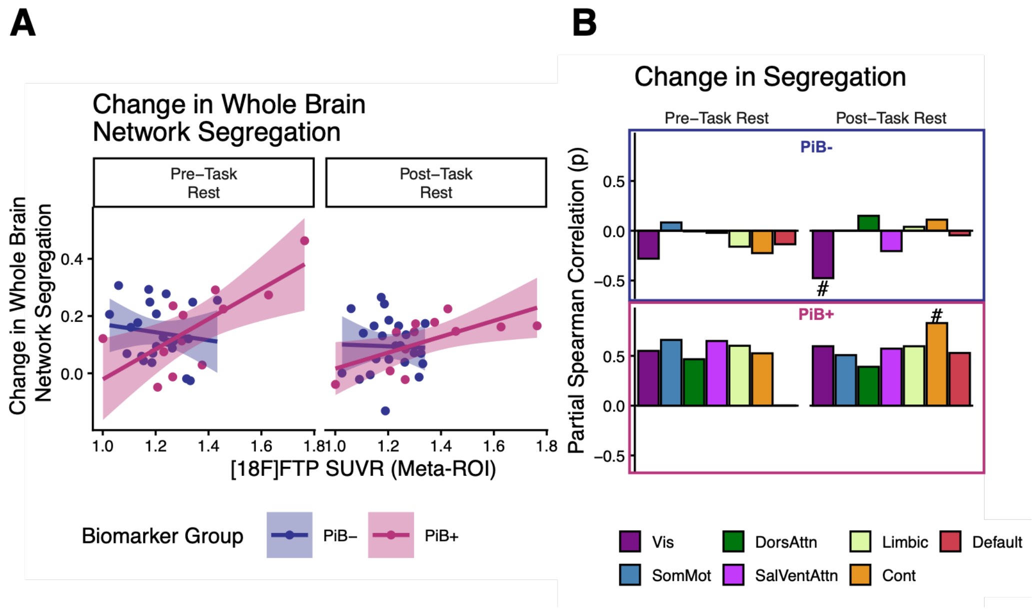


**Figure S4. Tau burden is associated with change in network segregation in PiB+ older adults. (A)** The relationship between tau burden in the meta-ROI ([$^{18}$F]FTP SUVR) and change in network segregation is plotted. A linear model testing for interactive effects of biomarker group and tau burden on change in network segregation, and controlling for age, sex, years of education, and head motion, found a significant interactive effect for change in network segregation calculated with pre-task rest (PiB+ vs. PiB-: $b = 0.87$, $t = 2.29$, $p = .03$). No interactive effect was found when change in network segregation was calculated with post-task rest (PiB+ vs. PiB-: $b = 0.61$, $t = 1.95$, $p = .06$). **(B)** Partial spearman correlation of change in network segregation (calculated separately with pre- and post-task rest) and tau burden is plotted separately for each Yeo-7 network. Age, sex, years of education, and head motion were partialed out. (# = $p < .05$ prior to FDR-correction; * = $q < .05$ after FDR-correction for multiple comparisons.)

**Figure S5.**

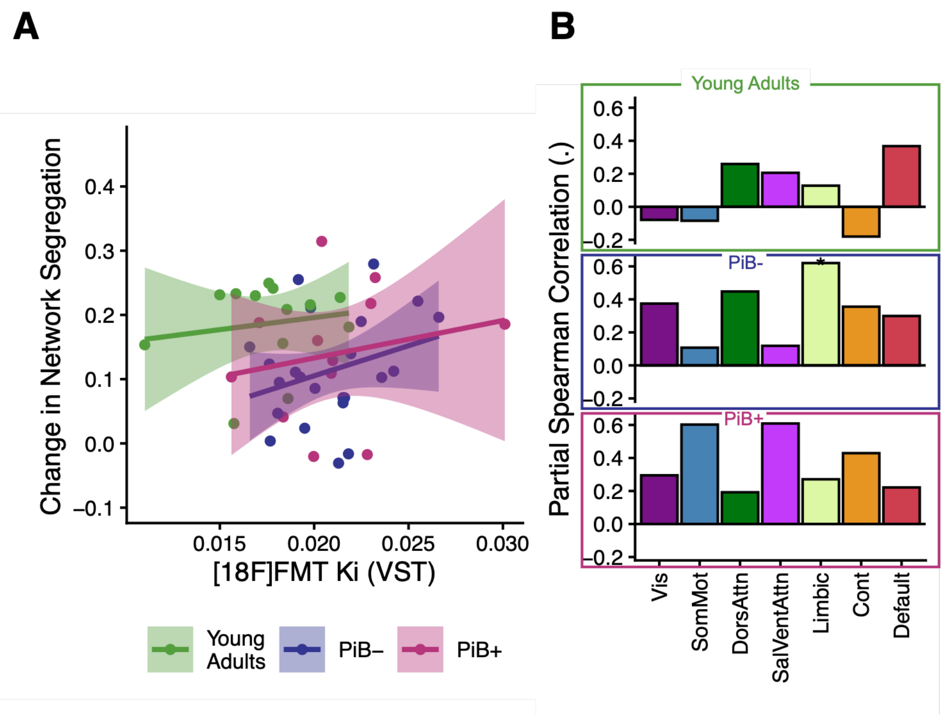


**Figure S5. (A)** Dopamine synthesis capacity was not significantly associated with changes in whole-brain network segregation at rest compared to task in any participant group. (Green = Young adults; Blue = PiB-; Pink = PiB+; Shaded region = 95% confidence interval). **(B)** In the PiB- group, there was a significant relationship between dopamine synthesis capacity and change in limbic network segregation from rest to task. Additionally, this relationship was detected for the PiB- group in the dorsal attention network, but was not significant after FDR correction for multiple comparisons. (Vis = Visual, SomMot = Somatomotor, DorsAttn = Dorsal Attention, SalVentAttn = Salience/Ventral Attention, Cont = Cognitive Control; # = $p < .05$, uncorrected; * = $q < .05$, FDR corrected for multiple comparisons).

**Figure S6.**

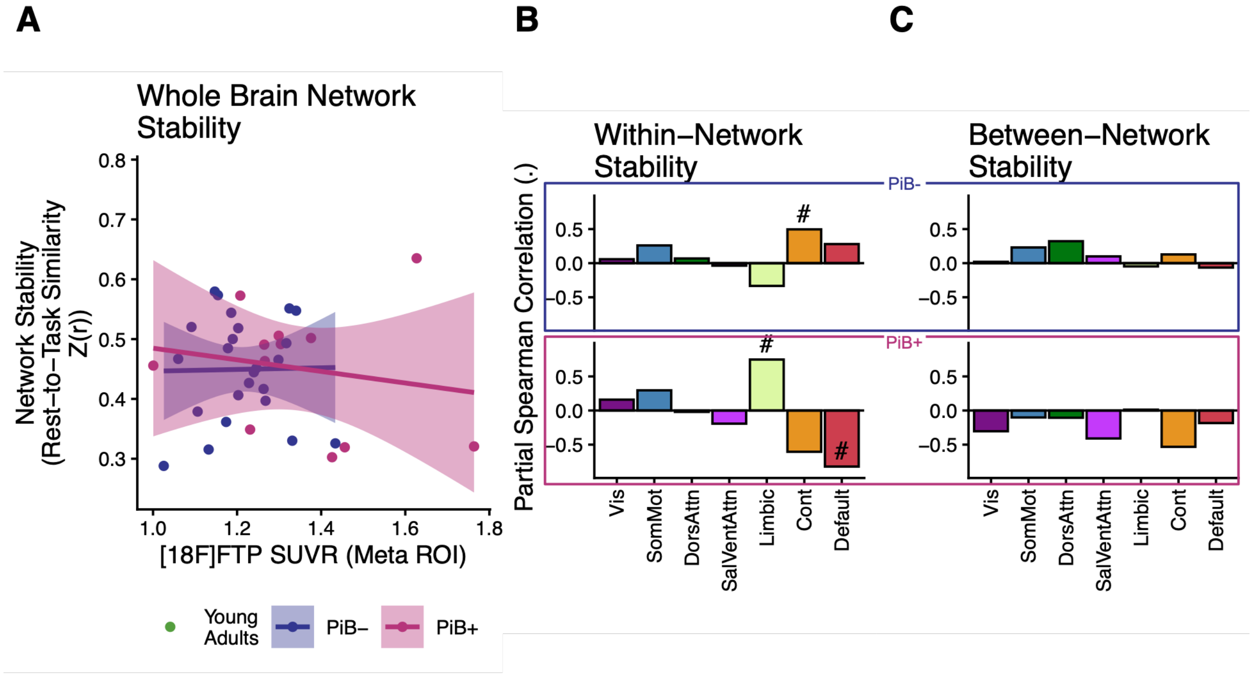


**Figure S6. Tau burden does not predict network stability in Older Adults. (A)** No significant relationship between tau burden and whole-brain network stability. **(B)** At the level of individual networks, stability within the cognitive control network was associated with tau burden in the PiB- group, and stability within the limbic/default mode networks was associated with tau burden in the PiB+ group, but these relationships did not survive correction for multiple comparisons. **(C)** Between-network stability was not significantly associated with tau burden in either group. (Trend lines = linear fit, Shaded area = 95% confidence interval. Green = Younger Adults, Blue = PiB-, Pink = PiB+; VST = ventral striatum, Vis = Visual, SomMot = Somatomotor, DorsAttn = Dorsal Attention, SalVentAttn = Salience/Ventral Attention, Cont = Cognitive Control; # = $p < .05$, uncorrected; * = $q < .05$, FDR corrected for multiple comparisons).
